# Molecular Basis of Cell Membrane Adaptation in Daptomycin-Resistant *Enterococcus faecalis*

**DOI:** 10.1101/2023.08.02.551704

**Authors:** April H. Nguyen, Truc T. Tran, Diana Panesso, Kara Hood, Vinathi Polamraju, Rutan Zhang, Ayesha Khan, William R. Miller, Eugenia Mileykovskaya, Yousif Shamoo, Libin Xu, Heidi Vitrac, Cesar A. Arias

## Abstract

Daptomycin is a last-resort lipopeptide antibiotic that disrupts cell membrane (CM) and peptidoglycan homeostasis. Enterococcus faecalis has developed a sophisticated mechanism to avoid daptomycin killing by re-distributing CM anionic phospholipids away from the septum. The CM changes are orchestrated by a three-component regulatory system, designated LiaFSR, with a possible contribution of cardiolipin synthase (Cls). However, the mechanism by which LiaFSR controls the CM response and the role of Cls are unknown. Here, we show that cardiolipin synthase activity is essential for anionic phospholipid redistribution and daptomycin resistance since deletion of the two genes (*cls1* and *cls2*) encoding Cls abolished CM remodeling. We identified LiaY, a transmembrane protein regulated by LiaFSR, as an important mediator of CM remodeling required for re-distribution of anionic phospholipid microdomains via interactions with Cls1. Together, our insights provide a mechanistic framework on the enterococcal response to cell envelope antibiotics that could be exploited therapeutically.

## Introduction

Enterococci are commensal bacteria that cause recalcitrant hospital-associated infections, and effective treatment has become a major challenge due to rising rates of antimicrobial resistance. Daptomycin (DAP) is a lipopeptide antibiotic with *in vitro* bactericidal activity against enterococci. While the mechanism of action of DAP is not fully understood, DAP is thought to bind to the cell membrane (CM) in a complex with anionic phospholipids (APLs) such as phosphatidylglycerol (PG) at the division septum^1–3^. DAP is then thought to insert into the membrane, where interaction with membrane lipid and protein components disrupts CM homeostasis and cell wall metabolism, ultimately leading to bacterial death^1,2^.

In *Enterococcus faecalis (Efs)*, DAP-resistance (DAP-R) is mediated by LiaFSR, a three-component cell envelope stress response system. Activation of the system causes CM changes, including alterations in lipid content and localization of APL microdomains-, a hallmark of DAP-R^3–7^. Previously, we discovered LiaX as an effector of the LiaFSR CM response with its C-terminal regulating the CM response against DAP through an unknown mechanism^4^.

DAP-R has also been associated with mutations in cardiolipin synthase (Cls), which synthesizes cardiolipin (CL) using PG as substrate^5,8,9^. CL content appears to affect several CM functions, including cell division^10–13^, and previous studies have linked increased levels of CL to DAP-R in *Efs* and *Staphylococcus aureus*^13–17^. *Efs* harbors two *cls* genes (*cls1* and *cls2*) with the majority of mutations associated with DAP-R identified in *cls1*^5,8,9^. Indeed, an R218Q substitution in the putative catalytic domain of Cls1 resulted in gain-of-function activity^15^. Experimental evolution of a DAP-susceptible (DAP-S) strain of *Efs* under DAP exposure has shown that Cls substitutions commonly arise following changes in LiaFSR, leading to high-level DAP-R^9^, suggesting that Cls may function in conjunction with LiaFSR to modulate CM adaptation. Here, we aimed to characterize the mechanistic basis of LiaFSR and Cls-mediated changes in CM adaptation that lead to DAP-R. We characterized a novel transmembrane protein (designated LiaY) that, with Cls, bridges the gap between sensing of DAP, alterations in phospholipid architecture and development of DAP-R.

## Results

### The *cls* genes have overlapping functions and are required for APL redistribution in DAP resistance

Both Cls1 and Cls2 synthesize CL^13^ yet, DAP-R mutations have been identified primarily in *cls1*^5,8,9^. Since Cls1 may function with the LiaFSR system to modulate the CM adaptive response^9^, we initially evaluated how activation of LiaFSR affects the expression of *cls1* and *cls2* using a DAP-R strain that lacks *liaX* (*Efs* OG117Δ*liaX*)^4,18^. Of note, deletion of *liaX* or the region coding for the C-terminus, constitutively activates the LiaFSR system and results in DAP-R^4^.

qRT-PCR^19^ showed that *cls1* expression was increased in DAP-R *Efs* OG117Δ*liaX* relative to its DAP-S parental strain (*Efs* OG117) (**Figure 1A**). In contrast, while *cls2* expression was increased in exponential phase growth, there was a sharp decrease in *cls2* expression in the stationary phase in the DAP-R strain (**Figure 1A**). These results suggest that the dynamics of *cls* gene expression differ between *cls1* and *cls2* within the cell cycle and support a predominant role for Cls1 in DAP-R. To test whether *cls1* and *cls2* were able to compensate for each other, we evaluated their expression in mutants containing individual deletions of either *cls1* or *cls2* in both *Efs* OG117 and *Efs* OG117Δ*liaX*. **Figure 1B** shows that, independent of the genetic background, deletion of either *cls* ultimately results in upregulation of the remaining *cls*, supporting functional redundancy between them (**Figure 1B**).

**Figure 1.**
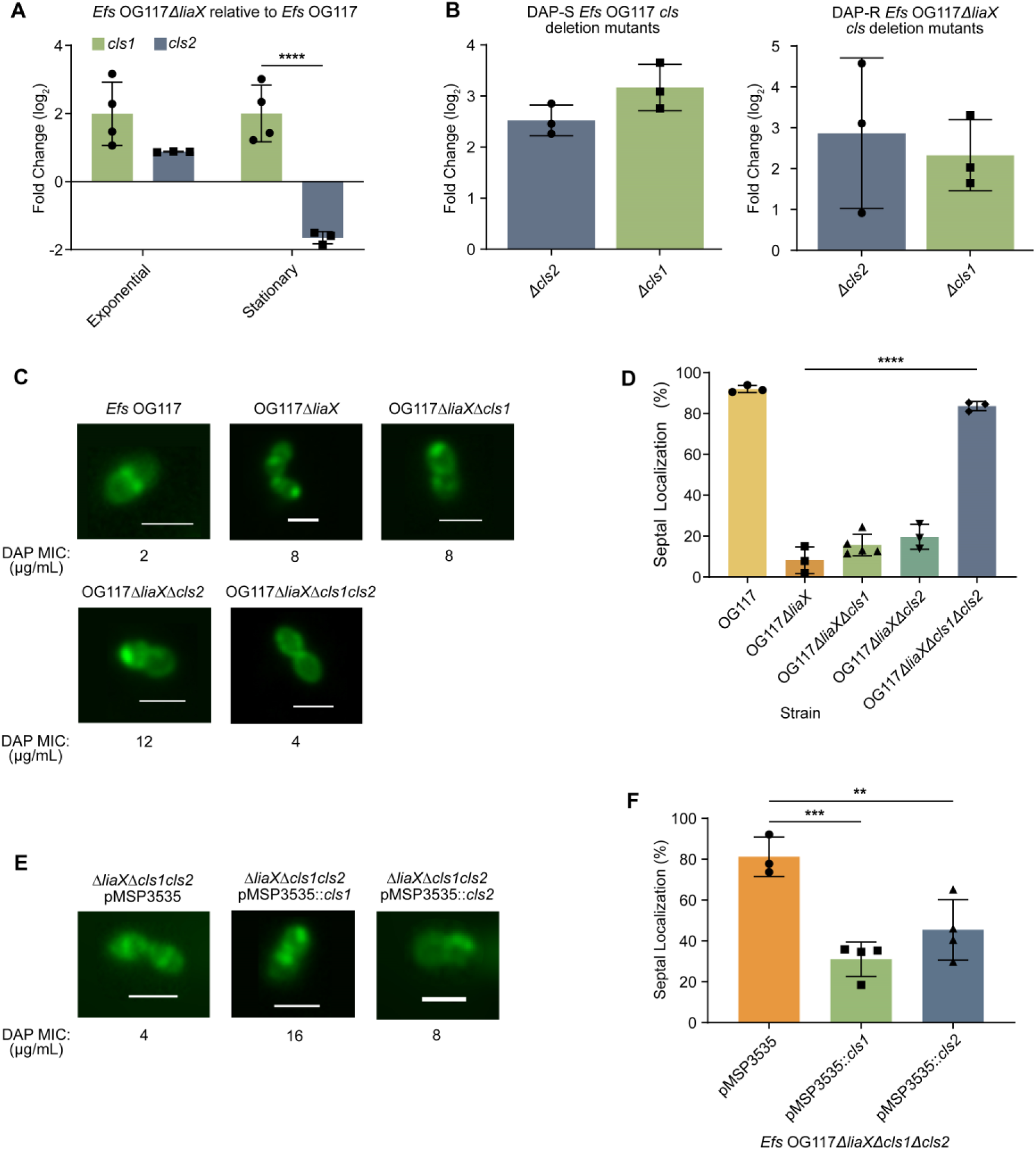
The *cls* genes have overlapping functions and are required for anionic phospholipid redistribution in DAP resistance. **(A)** Quantitative real time-PCR results evaluating *cls1* and *cls2* expression in DAP-R *Efs* OG117Δ*liaX* relative to DAP-S *Efs* OG117, ****p<0.0001, n=3-4 replicates **(B)** left panel, *cls* gene expression in *Efs* OG117Δ*cls1* and *Efs* OG117Δ*cls2* relative to *Efs* OG117; right panel, *cls* gene expression in *Efs* OG117Δ*liaX*Δ*cls1* or *Efs* OG117Δ*liaX*Δ*cls2* relative to *Efs* OG117Δ*liaX;* n=3 experiments **(C)** Representative images of NAO staining in *cls* mutant derivatives of *Efs* OG117Δ*liaX* and DAP MICs. Scale bar (white) at 2µm. **(D)** Quantification of septal localization of anionic phospholipid microdomains with NAO in *cls* mutant derivatives of *Efs* OG117Δ*liaX* by counting a minimum of 50 cells per replicate (n=3-5 replicates). **p<0.01, ***p<0.001, ****p<0.0001 **(E)** Representative images of NAO staining in *cls* mutant derivatives of *Efs* OG117Δ*liaX* complemented with vector pMSP3535 and derivatives **(F)** Quantification of septal localization of anionic phospholipid microdomains with NAO in *cls* mutants of *Efs* OG117Δ*liaX* complemented with vector pMSP3535 and derivatives by counting a minimum of 50 cells per replicate (n=3-4). **p<0.01, ***p<0.001, ****p<0.0001 Whole images were adjusted for “Black Balance” per BZ-X800 Image Analysis Software with individual representative selected.

Deletion of *cls1* or *cls2* individually did not alter DAP-associated phenotypes in the DAP-S *Efs* OG117. Indeed, DAP minimum inhibitory concentration (MIC) remained within susceptible levels (1-1.5 µg/mL) and APL microdomains exhibited “wild-type” septal localization, as visualized by 10-N-nonyl acridine orange (NAO) staining (**Figure S1A**). Similarly, individual deletions of either *cls1* or *cls2* in DAP-R OG117Δ*liaX* did not alter the DAP MIC or APL microdomain localization (non-septal pattern), with strains remaining DAP-R (8-12 µg/mL, **Figure 1C** and **Table S1**). These findings were confirmed through quantification of a subset of cells (n=50) to determine whether fluorescence intensity was concentrated at the septal or non-septal areas (**Figure 1D and Figure S1B**).

Subsequently, we generated tandem deletions of *cls1* and *cls2* to determine the impact of the complete absence of *cls* genes. Similar to single *cls* deletions, double deletions of both *cls* genes in DAP-S OG117 did not alter DAP MIC or APL localization (**Figure S1**). In contrast, a double deletion of both *cls1* and *cls2* in DAP-R *Efs* OG117Δ*liaX* reverted the resistance phenotype (**Table S1 and Figure 1C**) and restored septal localization of APL microdomains (**Figures 1C and 1D**).

To confirm the specific role of the *cls* genes in membrane adaptation, we trans-complemented the *Efs* OG117Δ*liaXΔcls1Δcls2* mutant with *cls1* and *cls2* genes individually, using a nisin-induced plasmid (pMSP3535)^20^. Expression of either *cls1* or *cls2* from pMSP3535 increased the DAP MICs and restored the non-septal APL microdomain localization (**Table S1**, **Figure 1E and 1F)**. We also cloned *cls1* and *cls2* independently in pAT392 under the constitutive promoter P2^21^. Constitutive expression of *cls1* in *Efs* OG117Δ*liaX*Δ*cls1*Δ*cls2* also increased the DAP MIC and reverted the septal localization of APL microdomains (**Fig S2A and S2B**). Of note, constitutive expression of *cls2* using pAT392 in *Efs* OG117Δ*liaXΔcls1Δcls2* resulted in a clumping phenotype with an unclear pattern of APL microdomain localization. However, DAP MIC was still increased (**Figure S2A**). Overall, our results suggest a major role for Cls1 in CM adaptation and indicates that CL is the major phospholipid species required for changes in membrane architecture that result in DAP-R.

### CM phospholipid changes associated with DAP-R and deletion of *cls* genes

To better understand the biochemical effects of *cls* deletions, we performed membrane lipid analysis using hydrophilic interaction liquid chromatography-ion mobility-mass spectrometry (HILIC-MS)^22,23^ on stationary phase cells, where changes in *cls* expression and CL content have been documented in the context of DAP-R^4^. As CL is a terminal product of a phospholipid biosynthetic pathway, changes in *cls* may trigger alterations in other lipid classes within the same or interrelated pathways (i.e. PG, lysyl-PG, diacylglycerol [DG], diglycodiacylglyercol [DGDG]^4,13,22,24,25^).

We first standardized lipid content of our representative DAP-S and DAP-R strains (*Efs* OG117 vs *Efs* OG117Δ*liaX*) under standard growth conditions (see Materials and Methods), depicting the lipid content as abundance of each lipid class per strain. The results indicate that, compared to DAP-S *Efs* OG117, DAP-R *Efs* OG117Δ*liaX* exhibited an increase in CL content, concomitant with a decrease in lysyl-PG and no significant difference in PG content (**Figure 2A**). We also identified the type and levels of individual CL species present within each strain by the length of fatty acid chain and degree of unsaturation. DAP-R OG117Δ*liaX* exhibited increased levels of CL species containing longer chain fatty acids and higher unsaturation compared to DAP-S OG117 strain (**Table 1**).

**Figure 2.**
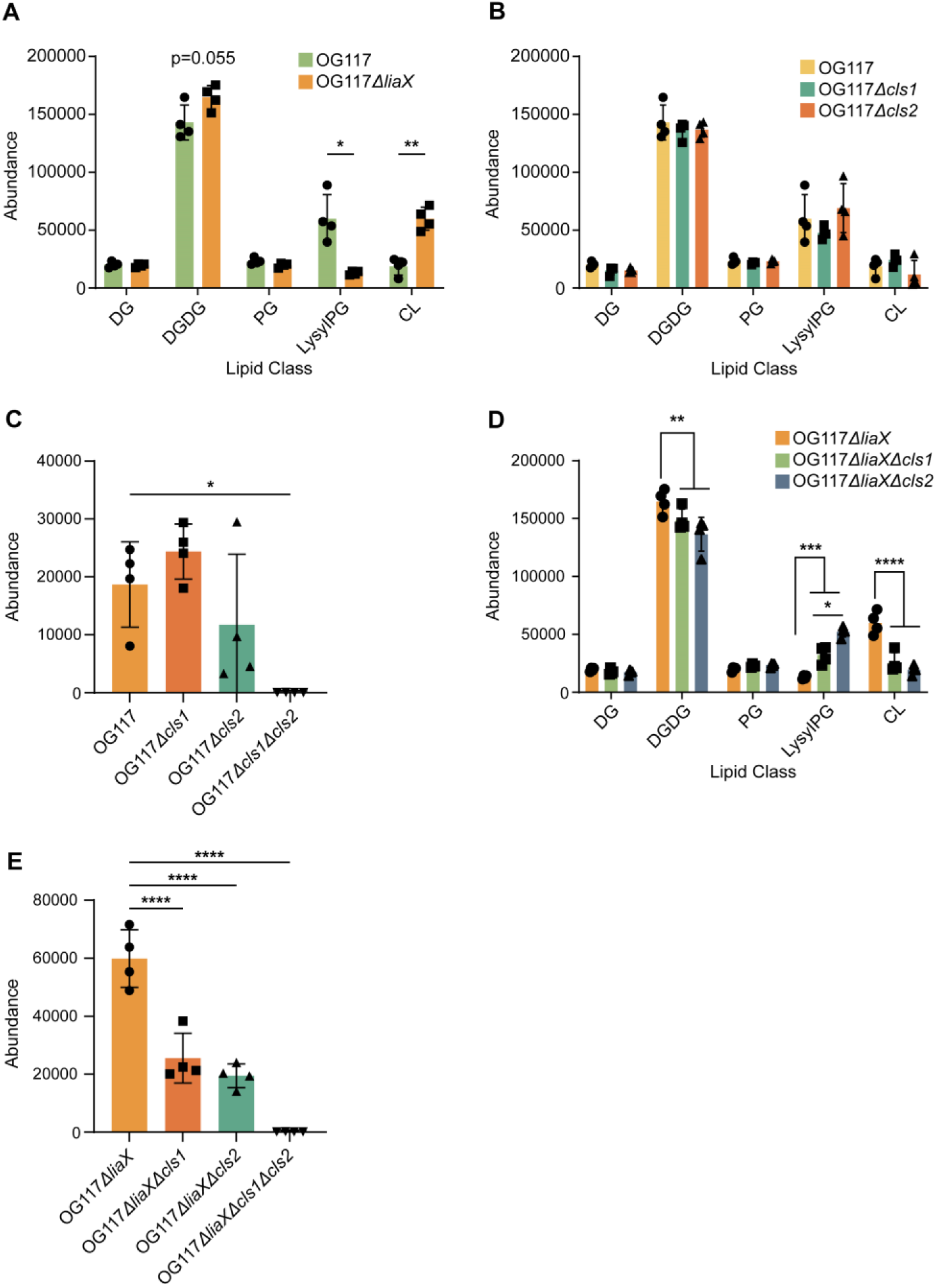
Cell membrane phospholipid changes associated with DAP-R and deletion of *cls* genes. **(A)** Quantification of lipid classes in DAP-susceptible *E. faecalis* OG117 and DAP-R *E. faecalis* OG117Δ*liaX* **(B)** Quantification of lipid classes in DAP-susceptible *E. faecalis*OG117 (parental) and *cls*mutant derivatives **(C)** Quantification of cardiolipin content in DAP-susceptible *E. faecalis* OG117 and *cls* mutant derivatives. **(D)** Quantification of lipid classes in DAP-R *E. faecalis* OG117Δ*liaX* and *cls* mutant derivatives. **(E)** Quantification of cardiolipin content in in DAP-R *E. faecalis* OG117Δ*liaX* and *cls* mutant derivatives. DG= diacylglycerol, DGDG = diglycodiacylglycerol, PG = phosphatidylglycerol, LysylPG = lysyl-phosphatidylglycerol, CL = cardiolipin *p<0.05 **p<0.01 ***p<0.001 ****p<0.0001, n=4

**Table 1.** Percentage of cardiolipin species present in *E. faecalis* OG117 (DAP-S) vs OG117Δ*liaX* (DAP-R)

| CL Species | OG117 (DAP-S) | OG117Δ*liaX* (DAP-R) |
| --- | --- | --- |
| CL 64:2 | **15.49 ± 1.40** | 8.96 ± 0.26 |
| CL 65:2 | **8.35 ± 0.47** | 4.92 ± 0.12 |
| CL 66:2 | **25.86 ± 0.77** | 22.21 ± 0.66 |
| CL 67:2 | **15.69 ± 0.71** | 13.00 ± 0.22 |
| CL 69:3 | 6.73 ± 0.23 | **7.94 ± 0.18** |
| CL 70:3 | 19.20 ± 1.19 | **26.79 ± 0.33** |
| CL 71:3 | 6.24 ± 0.75 | **9.75 ± 0.32** |
| CL 72:4 | 2.22 ± 0.59 | **5.41 ± 0.41** |
| CL 73:4 | 0.23 ± 0.14 | **1.03 ± 0.18** |

CL, cardiolipin; DAP, daptomycin. The first and second numbers (CL X:X) represent carbon chain length and number of double bonds (unsaturation), respectively. Table represents percentage of individual cardiolipin species identified from total amount of cardiolipin present. **Bold** text represents the strain with a statistically significant difference ( $p < 0.01$ ) in proportion of an individual CL species compared to the parental strain OG117. DAP-R OG117Δ *liaX* contains CL species with higher fatty acid chain length and unsaturation levels compared to DAP-S. The number represents the average of 4 independent experiments.

Individual deletions of either *cls1* or *cls2 in* DAP-S OG117 did not alter the overall content of CL or any other lipid classes tested compared to the parental strain. Importantly, there were also no differences in overall CL levels when comparing the *cls1* vs. *cls2* deletion in DAP-S OG117 (**Figure 2B**). Individual deletions of *cls1* or *cls2* caused a shift in CL species favoring fatty acids with longer chain length and higher degree of unsaturation compared to the parent *Efs* OG117 (**Table 2**). Of note, individual *cls* deletions showed no differences in CL species content compared to each other, further supporting the redundancy of their functions (**Table 2**). Most importantly, our lipidomics analyses confirmed that deleting both *cls1* and *cls2* completely abolished CL content in the membrane of DAP-S *Efs* OG117, demonstrating that Cls1 and Cls2 are true CL synthases^13^ (**Figure 2C**).

**Table 2.** Percentage of cardiolipin species present in *E. faecalis* OG117 (DAP-S) and *cls* mutant derivatives.

| CL Species | OG117 | OG117Δ*cls1* | OG117Δ*cls2* |
| --- | --- | --- | --- |
| CL 64:2 | **15.49 ± 1.4** | 12.6 ± 0.58 | 13.16 ± 1.38 |
| CL 65:2 | 8.35 ± 0.48 | 8.07 ± 0.61 | 7.16 ± 0.85 |
| CL 66:2 | **25.86 ± 0.77** | 22.05 ± 0.95 | 23.82 ± 1.16 |
| CL 67:2 | 15.69 ± 0.71 | 15.93 ± 0.52 | 15.54 ± 0.79 |
| CL 69:3 | **6.73 ± 0.23** | 8.17 ± 0.34 | 8.34 ± 0.5 |
| CL 70:3 | **19.2 ± 1.19** | 21.58 ± 0.73 | 21.97 ± 1.66 |
| CL 71:3 | **6.24 ± 0.76** | 7.77 ± 0.5 | 7.17 ± 1.32 |
| CL 72:4 | 2.22 ± 0.59 | 3.39 ± 0.37 | 2.6 ± 1.22 |
| CL 73:4 | 0.23 ± 0.14 | 0.44 ± 0.1 | 0.25 ± 0.22 |

CL, cardiolipin; DAP, daptomycin. The first and second numbers (CL X:X) represent carbon chain length and number of double bonds (unsaturation), respectively. The numbers represent the percentage of individual cardiolipin species identified from total amount of cardiolipin present. **Bold** text represents the strain with statistically significant ( $p < 0.05$ ) difference of each individual CL species comparing the parental strain with any of the *cls* mutants. The *cls* mutants show a shift towards fatty acids with higher fatty acid chain length and unsaturation levels. No significant differences between individual *cls* mutants were observed. The number represents the average of 4 independent experiments.

In contrast to the above findings, CM content analyses of DAP-R *Efs* OG117Δ*liaX* showed that individual *cls* deletions yielded a decrease in DGDG and CL content, as well as an increase in lysyl-PG content (**Figure 2D and 2E**). Furthermore, the overall amount of CL produced by *Efs* OG117Δ*liaX*Δ*cls1* or *Efs* OG117Δ*liaX*Δ*cls2* was very similar to that of DAP-S *Efs* OG117Δ*cls1* and *Efs* OG117Δ*cls2* (**Figure 2D**). However, the CL species produced by *Efs* OG117Δ*liaX*Δ*cls1* and *Efs* OG117Δ*liaX*Δ*cls2* harbored shorter chain fatty acids and lower degree of unsaturation when compared to the parent *Efs* OG117Δ*liaX* **(Table 3).** Nonetheless, when comparing individual *cls* deletions between each other in this DAP-R background, we found no statistically significant differences in levels of CL species except for CL 64:2 and 70:3 (**Table 3**). Finally, similar to OG117, deletion of both *cls1* and *cls2* in DAP-R *Efs* OG117Δ*liaX* resulted in total abolishment of CM CL content (**Figure 2E**).

**Table 3.** Percentage of cardiolipin species present in *E. faecalis* OG117Δ*liaX* (DAP-R) and *cls* mutant derivatives.

| CL Species | OG117Δ*liaX* | OG117Δ*liaX*Δ*cls1* | OG117Δ*liaX*Δ*cls2* |
| --- | --- | --- | --- |
| CL 64:2 | **8.96 ± 0.26** | 10.74 ± 0.42* | 13.35 ± 0.57 |
| CL 65:2 | 4.92 ± 0.12 | 4.3 ± 0.28 | 5.29 ± 0.28 |
| CL 66:2 | **22.21 ± 0.66** | 25.78 ± 0.45 | 26.8 ± 0.6 |
| CL 67:2 | 13.01 ± 0.22 | 12.43 ± 0.42 | 13.02 ± 0.16 |
| CL 69:3 | **7.94 ± 0.18** | 7.03 ± 0.14 | 7.04 ± 0.1 |
| CL 70:3 | 26.79 ± 0.33 | 26.8 ± 0.78* | 23.49 ± 0.51 |
| CL 71:3 | **9.75 ± 0.32** | 8.72 ± 0.32 | 7.57 ± 0.38 |
| CL 72:4 | **5.41 ± 0.41** | 3.81 ± 0.37 | 3.16 ± 0.27 |
| CL 73:4 | 1.03 ± 0.18 | 0.41 ± 0.14 | 0.29 ± 0.09 |

CL, cardiolipin; DAP, daptomycin. The first and second numbers (CL X:X) represent carbon chain length and number of double bonds (unsaturation), respectively. Table represents percentage of individual cardiolipin species identified from total amount of cardiolipin present. Text in **bold** represents a statistically significant ( $p < 0.05$ ) difference in the proportion of each individual CL species from the *cls* mutants with the parental strain OG117Δ*liaX*. \*Represents a statistically significant difference ( $*p < 0.05$ ) in a CL species between the individual *cls* mutants (CL 64:2, 70:3). Overall, the *cls* mutants show shift towards fatty acids with shorter fatty acid chain length and unsaturation levels.

Put together, our results suggest that CL plays a major role in DAP-R and that the phenotype results in increased CL content with longer fatty acids and a higher degree of unsaturation. This biochemical signature appears to be mediated by contributions and overlapping functions of both Cls1 and Cls2.

### Cls1 localizes in non-septal APL microdomains in DAP-R *Efs*

Our initial results provided evidence that the formation of non-septal anionic phospholipid microdomains is dependent on the availability of CL and Cls. We hypothesized that Cls1 is likely responsible for the formation of non-septally located APL microdomains. Therefore, we evaluated the membrane localization of a tetracysteine-tagged Cls1 by fluorescence microscopy using a ReAsH biarsenical reagent^26^ coupled with NAO staining in both in DAP-S *Efs* OG1RF and DAP-R *Efs* OG1RFΔ*liaX.* **Figure 3A** shows that, **i)** in DAP-S *Efs* OG1RF, Cls1 is localized at the septum, **ii)** in DAP-R *Efs* OG1RFΔ*liaX,* Cls1 is localized non-septally and, **iii)** Cls1 was co-localized with APL microdomains in both conditions (**Figure 3A**). To confirm the co-localization of Cls1 and APL microdomains, we cloned the gene expressing ReAsH-Cls1 in pMSP3535 and transformed it into OG1RFΔ*liaX*Δ*cls1*Δ*cls2*. **Figure 3B** shows that, in the absence of CL, APL microdomains are mainly visualized at the septum. Introduction of *reAsH-cls1* restored the non-septal localization of the APL microdomains, confirming its essentiality in mediating CM remodeling in DAP-R. Furthermore, since we previously showed that Cls1 and Cls2 have redundant functions in DAP-R (see above), we also investigated the localization dynamics of a GFP-tagged Cls2 in both DAP-S and DAP-R with strains lacking *cls1* showing septal localization of Cls2 in DAP-S *Efs* OG117Δ*cls1* and non-septal localization of Cls2 in DAP-R OG117Δ*liaX*Δ*cls1* **(Figure S3**). Taken together, our results highlight the major role of Cls in triggering changes in membrane architecture mediating DAP-R.

**Figure 3.**
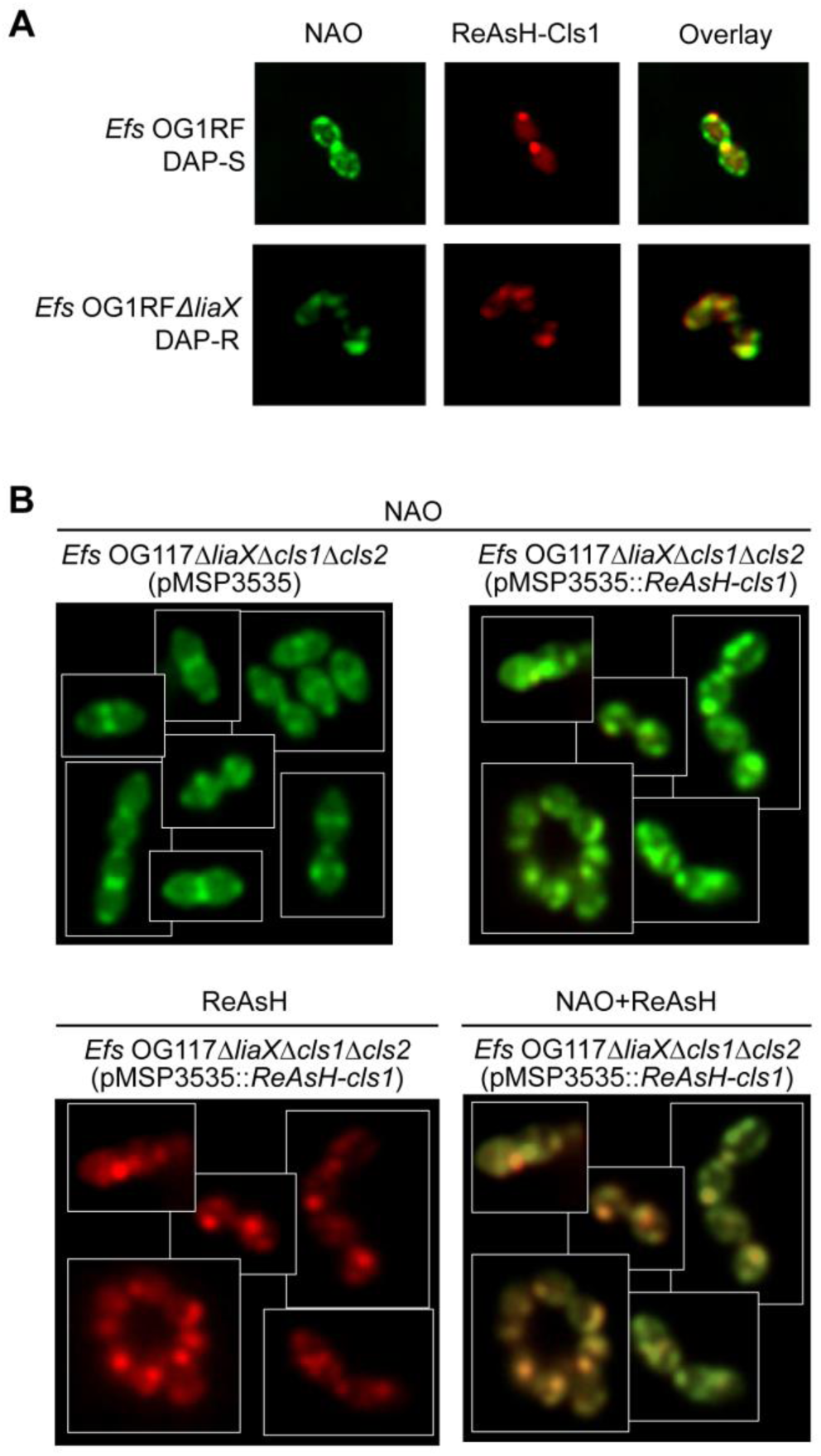
Cls1 localizes in non-septal anionic phospholipid microdomains in DAP-R *E. faecalis*. **(A)** Representative images of 10-N-nonyl acridine orange (NAO) staining (left panel), tetracysteine-tagged Cls1 with ReAsH reagent (red fluorescence, middle panel) and overlay of both images (right panel). Both anionic phospholipid microdomains and Cls1 are shown to relocalize and overlap away from the septum in DAP-R *Efs* (OG1RFΔ*liaX*) (Pearson Correlation Coefficient 0.91, [95% CI: 0.74, 0.97]) compared to their septal pattern in DAP-S *E. faecalis* OG1RF (Pearson Correlation Coefficient 0.94, [95% CI; 0.83, 0.98]). **(B)** NAO staining for anionic phospholipid microdomains (green) and tetracysteine tagged-Cls1 localization (red) in *E. faecalis* OG117Δ*liaX*Δ*cls1*Δ*cls2* harboring pMSP3535 or pMSP3535 carrying the gene expressing the tetracysteine tagged-Cls1. Right far panel, overlap between NAO and ReAsH (Pearson Correlation Coefficient 0.91, [95% CI: 0.76, 0.96]) Whole images were adjusted for “Black Balance” per BZ-X800 Image Analysis Software with individual representative selected.

### LiaY, a member of the LiaFSR system, mediates CM remodeling associated with DAP-resistance

The main target of the LiaR response regulator is a three-gene cluster designated *liaXYZ*. We previously showed that LiaX plays a major role in controlling the activation of the CM response through its C-terminal domain^4^. Here, we sought to characterize the role of *liaYZ* in DAP-associated CM remodeling. *In silico* evaluation indicated that LiaY is a 107 amino acid transmembrane protein that is predicted to contain a PspC domain (involved in the phage shock protein stress response in *E. coli*)^27^. LiaZ is a 118 amino acid transmembrane protein that shows homology with holins, generally involved in autolysis^28^. We initially attempted to generate knockout mutants^29,30^ of *liaY*, *liaZ* and both genes in the background of DAP-R *Efs* OG1RF *liaX*289* (harboring a deletion of the region encoding the C-terminus of LiaX^4^). In *Efs* OG1RF *liaX*289*, deletion of *liaZ* had a marginal effect on DAP MICs (decreasing 1.5 fold) and predominantly exhibiting a non-septal pattern of APL microdomains (**Figures 4A and 4B**). We were unable to delete *liaY* in *Efs* OG1RF *liaX*289* without generating compensatory mutations in *liaS* (see Discussion). In contrast, we successfully generated individual *liaY* mutants in DAP-S OG1RF without additional mutations in *liaFSR*. Of note, characterization of these *liaY* mutant strains indicated that deletion of *liaY* in DAP-S OG1RF (without an activated LiaFSR system) did not affect the DAP MIC or APL microdomain architecture (**Figure S4**), supporting the notion that LiaY is critical when the LiaFSR is activated to maintain the DAP-R phenotype.

**Figure 4.**
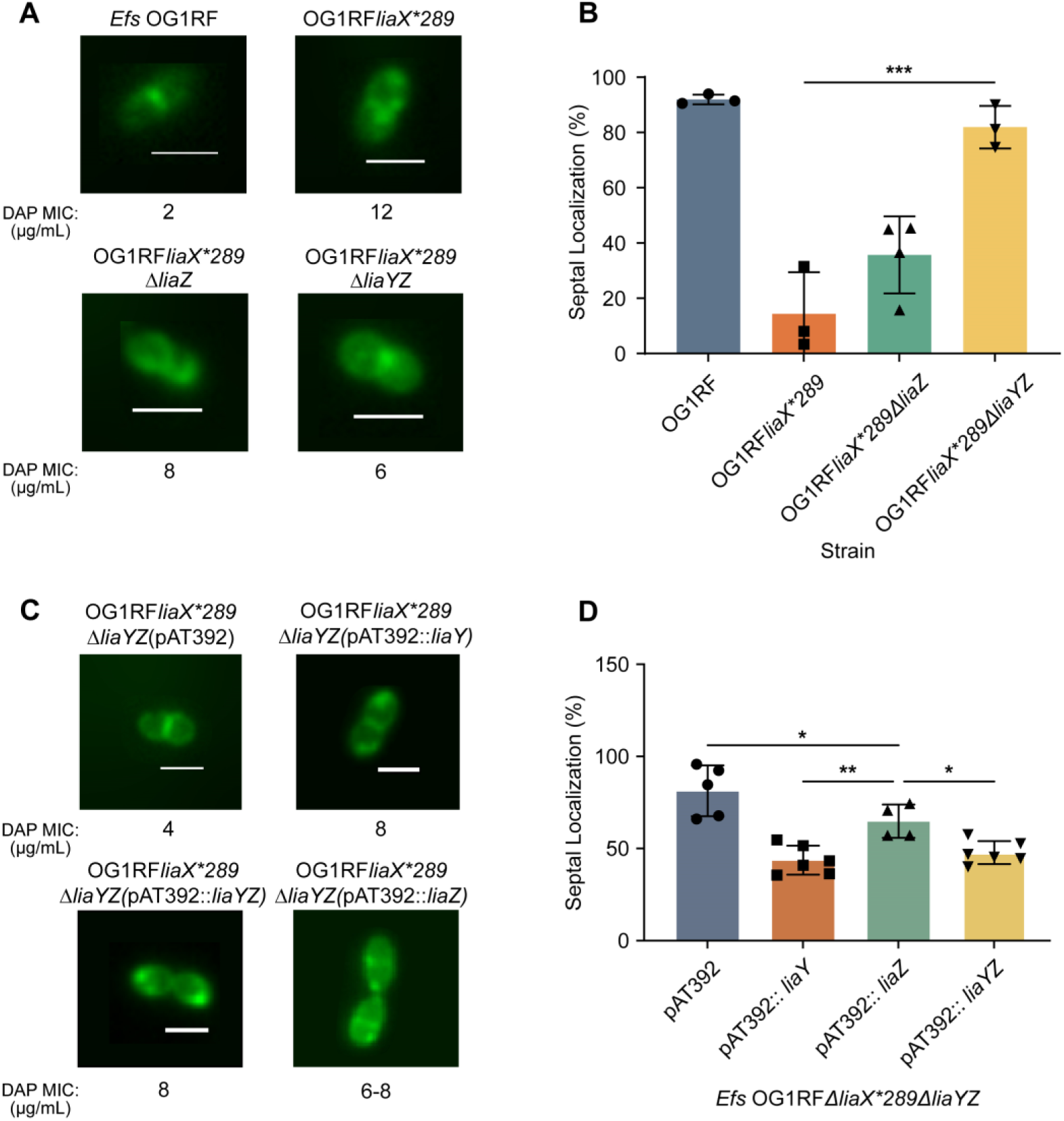
LiaY, a member of the LiaFSR system, mediates changes in phospholipid architecture associated with DAP-resistance. **(A)** Representative images of anionic phospholipid localization by NAO staining in DAP-R OG1RFΔ*liaX*289*, and *liaXYZ* mutants. Scale bar (white) at 2 µM. **(B)** Quantification of septal localization of anionic phospholipid microdomains by NAO staining in OG1RF and *liaXYZ* mutants (>50 cells counted per strain per replicate, n=3-4 replicates). *p<0.05; **p<0.001; ***p<0.001. **C)** Representative images of anionic phospholipid localization by NAO staining in OG1RF *liaX*289*Δ*liaYZ* mutant transformed with pAT392 and derivatives. Scale bar (white) at 2µM. **D)** Quantification of septal localization of anionic phospholipid microdomains by NAO staining of OG1RF *liaX*289*Δ*liaYZ* mutant transformed with pAT392 and derivatives (>50 cells counted per strain per replicate, n=6-9 replicates). *p<0.05; **p<0.001; ***p<0.001;****p<0.0001. Whole images were adjusted for “Black Balance” per BZ-X800 Image Analysis Software with individual representative selected.

Despite difficulties in generating *liaY* mutants in DAP-R *Efs* OG1RF *liaX*289*, we were able to make a double mutant of *liaYZ*. The double mutant had a reduced DAP MIC (6 μg/mL compared to 12 μg/mL in the parent strain) (**Table S1**) with restoration of septal APL microdomain localization (**Figures 4A and 4B**). These findings are consistent with our hypothesis that *liaY* was important in the generation of non-septal phospholipid microdomains associated with DAP-R.

To confirm the role of *liaY* in CM remodeling without specifically generating a *liaY* deletion, we opted to introduce *liaY, liaZ, or liaYZ* into the double mutant *Efs* OG1RF *liaX*289*Δ*liaYZ* using pMSP3535 (nisin inducible)^20^ and pAT392^21^. Trans-complementation of *Efs* OG1RF *liaX*289*Δ*liaYZ* with *liaY* on pMSP3535 **(Figure S2C and S2D)** or pAT392 **(Figure 4C and D)** caused a marked increase in DAP MICs in OG1RF *liaX*289*Δ*liaYZ* (3 and 2-fold, respectively) (**Table S1**). We also identified an increase in the non-septal localization of APL microdomains, confirming the role of LiaY in DAP-mediated CM adaptation. Constitutive expression of *liaZ* in *Efs* OG1RF *liaX*289*Δ*liaYZ* also caused an increase in DAP MIC. However, the effect on APL microdomain localization was less pronounced compared to *trans* expression of *liaY* or *liaYZ* **(Figure 4C and D, Table S1)**. Of note, introduction of *liaZ* under the constitutive promoter P2 in pAT392 resulted in a proportion of the population with a clumping phenotype similar to that observed with *cls2* expression in OG117Δ*liaX*Δ*cls1*Δ*cls2* (**Figure S2A and S5**). Thus, our results indicate that both LiaY and LiaZ contribute to the redistribution of APL microdomains associated with DAP-R, although LiaY seems to play the more dominant role in this effect.

### LiaY bridges the LiaFSR response with changes in membrane architecture via Cls

Since both Cls and LiaY are crucial for the formation of non-septal APL microdomains associated with DAP-R, we investigated if this adaptation was mediated by interactions of LiaY with Cls1 and evaluated this possibility by applying the bacterial two-hybrid system as an initial screen. LiaY and Cls were tagged either at the N- or C-terminus of adenylate cyclase reporter domains with beta-galactosidase activity as a measure of potential interaction. **Figure 5A** shows that the LiaY and Cls1 have a likely interaction. Of note, we were not able to detect a potential interaction between Cls1 and LiaZ (**Figure 5B**), confirming that LiaY was the main member of the LiaFSR system involved in membrane remodeling likely via interactions with Cls1.

**Figure 5.**
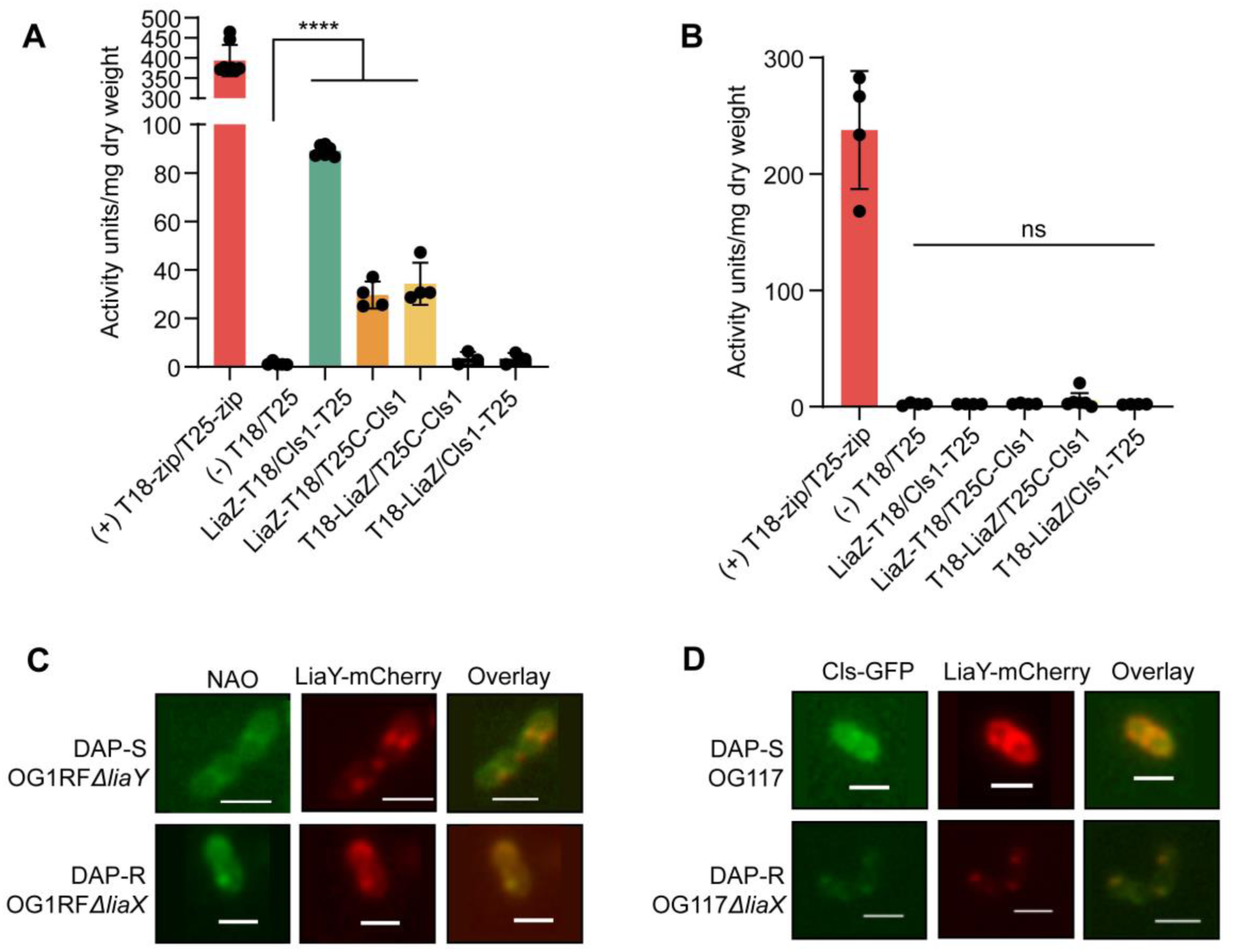
LiaY bridges the LiaFSR response with changes in membrane architecture via Cls. **(A)** Protein-protein interactions between LiaY and Cls1 using the bacterial two hybrid system. Proteins were tagged at either the N- or C-terminus, co-transformed into *E. coli* BTH101 and activity recorded via a beta galactosidase assay. A leucine zipper interaction was used as the positive control (T18-zip/T25-zip, red bar) with two non-tagged empty vectors used as negative controls (T18/T25). n= 3-8. *p<0.05; **p<0.001; ***p<0.001. **(B)** Protein-protein interactions between LiaZ and Cls1 using the bacterial two hybrid system and following the same methodology as in (A). Ns, non-significant. n=4-7. *p<0.05; **p<0.001; ***p<0.001. **(C)** Representative images of NAO staining for anionic phospholipid microdomains (left panel), LiaY-mCherry (central panel) and overlay of both images (right panel) in DAP-S lacking *liaY* (*Efs* OG117Δ*liaY;* top panels) (Pearson Correlation Coefficient 0.67, [95% CI: 0.24, 0.88]) and DAP-R (*Efs* OG117Δ*liaX;* bottom panels) (Pearson Correlation Coefficient 0.91, [95% CI: 0.75, 0.97]). Both strains strains were transformed with pMSP3535::*liaY-mCherry*. **(D)** Co-localization of LiaY-mCherry and Cls1-GFP via fluorescence microscopy in *Efs* OG117 (Pearson Correlation Coefficient 0.69, [95% CI: 0.28, 0.89]) and *Efs* OG117Δ*liaX* GFP-Cls1 (Pearson Correlation Coefficient 0.86, [95% CI: 0.63, 0.95]) (with *gfp-cls1* introduced in the chromosome) transformed with pMSP3535::*mCherry-liaY*. Whole images were adjusted for “Black Balance” per BZ-X800 Image Analysis Software with individual representative selected.

We next used fluorescence microscopy to determine whether Cls1 and LiaY co-localize with each other and within APL microdomains either at the septum or away from the septum in DAP-S and DAP-R *Efs* strains, respectively. We generated a *liaY*-mCherry construct in pMSP3535 and were able to identify LiaY localized at the septum in DAP-S *Efs* OG1RFΔ*liaY* (**Figure 5C**). In contrast, LiaY was visualized non-septally in DAP-R *Efs* OG117Δ*liaX*. Most importantly, we were able to show that, similar to Cls1, LiaY co-localized to APL microdomains in both DAP-S *Efs* OG1RFΔ*liaY* and DAP-R *Efs* OG1RFΔ*liaX* (**Figure 5C)**. Of note, *Efs* OG1RFΔ*liaX* expressed both the native *liaY* and *liaY-mCherry* due to the difficulties of generating a *liaY* deletion mutant in DAP-R backgrounds.

Lastly, to confirm the LiaY-Cls1 co-localization, we next evaluated the *in vivo* interaction of Cls1 and LiaY using a GFP-tagged Cls1 introduced into the chromosome, concomitant with *liaY*-*mCherry* expressed on pMSP3535. **Figure 5D** shows that Cls1 and LiaY colocalize at the septum in DAP-S *Efs* OG117 and at non-septal sites in DAP-R *Efs* OG117Δ*liaX*. Controls for potential self-aggregation of fluorescent tags were included to rule out artifactual localization patterns using pMSP3535 expressing GFP or mCherry alone, showing only a diffuse pattern with no discrete foci in either *Efs* OG117 (DAP-S) or OG117Δ*liaX* (DAP-R) (**Figure S6**). Furthermore, quantification of colocalization between LiaY and NAO or LiaY and Cls1 showed much stronger association in DAP-R over DAP-S strains (**Figure 5C and 5D**), supporting our hypothesis that LiaY and Cls1 function together to mediate DAP-R when LiaFSR is activated.

## Discussion

DAP is thought to target fluid areas of the membrane rich in PG^1^ with eventual CM penetration likely causing inhibition of cell wall synthesis^2^ and compromised cell envelope integrity^1^. Enterococci counteract these negative effects by triggering a membrane adaptive response coordinated by members of the LiaFSR system^4,5,7,8^ to re-localize APL microdomains^6,31^, ultimately diverting DAP from the septum^4,6^. However, the mechanistic basis of this response are unknown. Here, we identified the major elements involved in this response and provide a mechanistic model to gain further understanding on how multidrug-resistant enterococci adapt their CMs to survive^32,33^.

We first characterized the role of CL and Cls, which have frequently been associated with DAP-R without a clear mechanism^5,8,9^. Our findings support a model in which Cls1 is upregulated upon development of DAP-R, acting as a bridge between activation of the LiaFSR system and DAP-R-associated changes in the CM. Interestingly, Cls1 is not regulated by LiaR^4^, and we speculate that changes in phospholipid metabolism upon oligomerization of DAP in the CM could trigger the increased expression of *cls1*. In any scenario, CL-rich, non-septal APL microdomains are likely to alter the CM oligomerization and translocation properties of DAP, as some previous published work suggests^14^, and “trap” DAP in areas away from the division septum to protect septal cell division and peptidoglycan synthesis^6^ (**Figure 6**).

**Figure 6.**
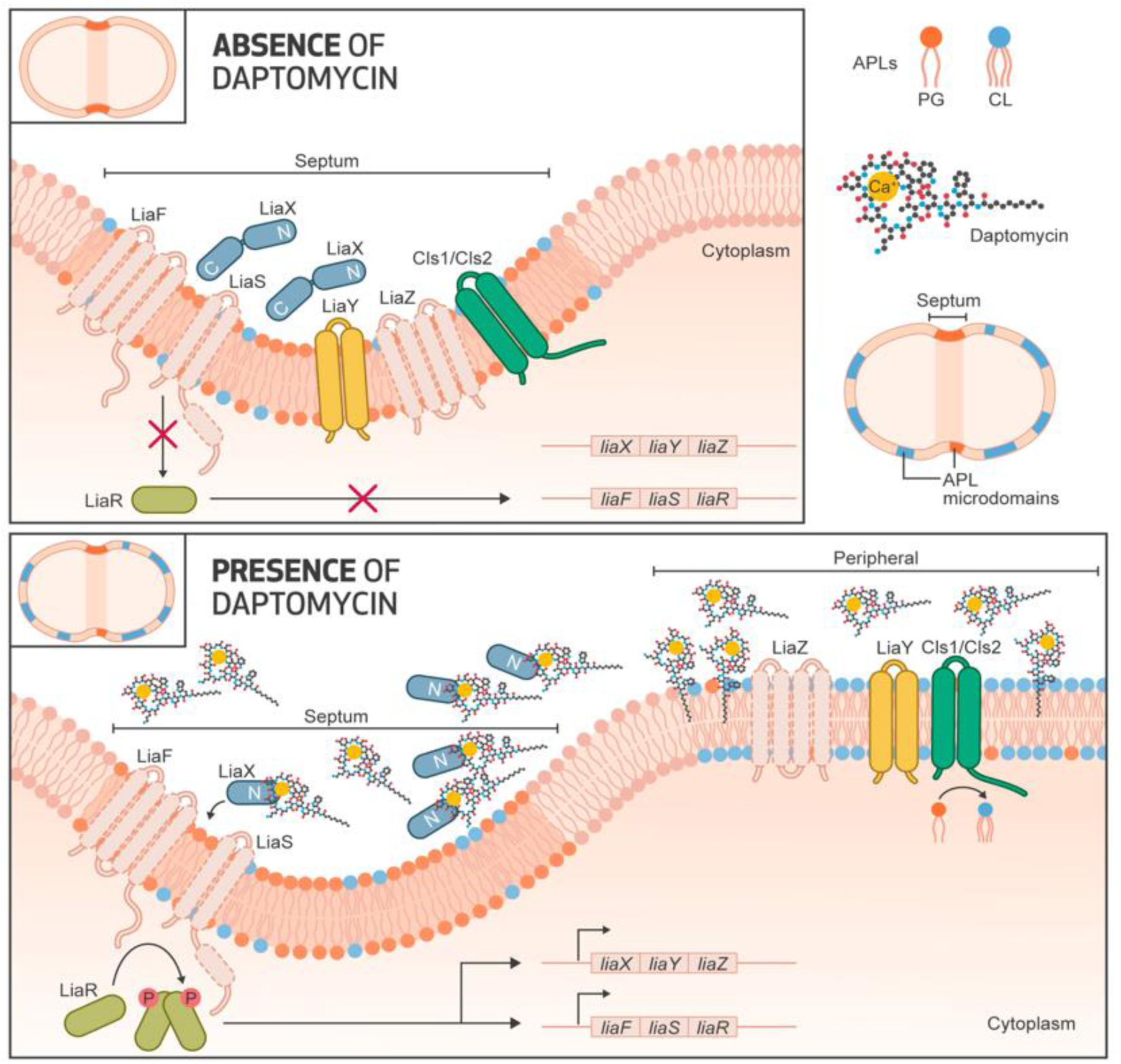
Mechanistic model of cell membrane adaptation in daptomycin-resistant *Enterococcus faecalis*. In the absence of daptomycin (**top panel**), anionic phospholipid microdomains are located at the bacterial division septum where other critical membrane-associated proteins involved in cell division, cell wall or lipid biosynthesis (such as Cls1) are also found. Members of the LiaFSR system (namely *liaXYZ*) are produced at basal levels and localize in septal areas. LiaX plays an inhibitory role through its C-terminal domain likely via interactions with members of the LiaFSR system. Upon activation of LiaFSR in the presence of daptomycin (**bottom panel**), the N-terminal domain of LiaX is released to the milieu and interacts with DAP. Concomitant changes in the C-terminus of LiaX cause “activation” of LiaY, generating interactions with Cls1. The LiaY-Cls1 complexes localize in areas away from the septum, presumably in regions where the membrane is being damaged by the antibiotic. Cls1 then produces high amounts of cardiolipin in these non-septal areas, attracting DAP molecules to these membrane regions resulting in alteration of the interaction of DAP with its transmembrane target (likely lipid II intermediates), preventing further DAP-mediated damage.

*Efs* possesses more than one copy of the *cls* gene, and our results showed that both Cls1 and Cls2 have overlapping functions. Although this functional redundancy is not surprising, the reasons for the preference for Cls1 during development of DAP-R are obscure as are any differential roles between the two enzymes. Of note, overexpression of *cls2* appears to have deleterious effects on cell division (**Figure S2A**), suggesting that Cls2 may have unique roles in certain biological contexts. Also, our lipidomics studies show a difference in levels of certain CL species synthesized by Cls1 or Cls2 (**Table 3**). These findings suggest that PG units containing particular fatty acids can be preferentially incorporated by individual Cls enzymes. Fatty acid tail length may allow CL species to alter the physicochemical properties of the CM, with previous studies showing that areas containing increased cyclic or decreased unsaturated fatty acids are likely less prone to the deleterious effects of DAP^24^. Our results also provide further insights onto the controversy surrounding the use of NAO staining. Historically, NAO has been proposed to bind CM regions that are rich in CL^34^. However, it has been shown that NAO binds other APLs, such as PG or phosphatidic acid, in conditions of low (or no) CL content^35^, and we also show positive NAO staining in the absence of detectable levels of CL.

Another major finding of this work is the discovery of the function of LiaY, a protein that bridges the LiaFSR system with phospholipid homeostasis via interactions with Cls. Interestingly, we could not generate a *liaY* mutant in DAP-R derivatives such as OG117Δ*liaX*, where the LiaFSR system is constitutively activated^4^. In DAP-R OG117Δ*liaX,* attempts to delete *liaY* consistently resulted in a concomitant mutation in *liaFSR* that, presumably, shuts off the LiaFSR response, and the strain became hypersusceptible to DAP. The *liaY* gene is located in a cluster of 3 genes (*liaXYZ*), previously shown to be regulated by LiaR^4^. Of note, *liaY* and the start codon of *liaZ* overlap, suggesting that they are co-transcribed. The inability to delete *liaY* in the DAP-R strain suggests that, under conditions of envelope stress, a stoichiometric balance between LiaY and LiaZ may influence cell viability in the absence of compensatory mutations. Supporting a deleterious role of unregulated LiaZ, plasmid expression of *liaZ* in *Efs* OG1RF *liaX*289*Δ*liaYZ* led to an abnormal “clumping” phenotype (**Figure S5**). Therefore, it is tempting to speculate that LiaY may modulate LiaZ function, and our studies showed a positive interaction between LiaY and LiaZ (**Figure S7**). In any case, our results indicate that both LiaY and Cls are both necessary and essential for the formation of non-septal anionic phospholipid microdomains.

Finally, we propose a mechanistic model of CM activation triggered by DAP that leads to resistance **(Figure 6)**. In DAP-S strains, APL microdomains are located at the bacterial division septum, favoring optimal functioning of critical proteins involved in vital cell processes, including cell division^10,11^ at the septum. In the absence of DAP, the LiaFSR system (and LiaXYZ) are produced at basal levels and localize in septal areas. Upon activation of LiaFSR, the increased expression of *liaXYZ* results in ***i)*** changes in LiaX^4^ that result in the “activation” of LiaY generating interactions with Cls1, ***ii***) the LiaY-Cls1 complexes localizing away from the septum, and ***iii***) Cls1 producing CL in these non-septal areas, preventing further DAP-mediated damage by altering the interaction of DAP with its septal targets. Moreover, the APL microdomains are likely to serve as a signal for aberrant cell division and attract proteins involved in the divisome (as it has been previously shown^10,11^), preventing damage of cell division processes at the septum.

Overall, our results provide evidence to bridge the activation of the LiaFSR system with CL homeostasis as an adaptive response at the membrane level leading to DAP-R and provide further insights onto the strategies that Gram-positive bacteria deploy to avoid the killing effect of cell envelope targeting antibiotics.

## Materials and Methods

### Materials Availability

Requests for use of specific generated strains can be directed to the lead contact and may be subject to a material transfer agreement.

### Data and Code Availability

Data reported in this paper will be shared by the lead contact upon request. This paper does no report original code. Microbial strains and microbial source material for *in vitro* studies are reported in **Table S1**. Culture conditions can be found in “Method Details.”

### Bacterial Growth

*E. coli* strains were cultured at 37°C with agitation either in broth or on agar plates with Luria-Bertani (Becton-Dickinson) with appropriate antibiotics (Sigma-Aldrich) added for maintaining plasmids if needed: 300 µg/mL erythromycin for pMSP3535^22^, 25 µg/mL gentamicin for pHOU1^31,32^/pAT392^23^, 50 µg/mL kanamycin or 100 μg ampicillin for bacterial two hybrid plasmids (see below), 15 µg/mL chloramphenicol for pCE^18^. Enterococcal strains were cultured at 37°C with agitation either in broth or on agar plates with Brain Heart Infusion or Tryptic Soy Broth (Becton-Dickinson) with appropriate antibiotics (Sigma-Aldrich) added for maintaining plasmids if needed: 15 µg/mL erythromycin for pMSP3535, 150 µg/mL gentamicin for pHOU1/pAT392, 15 µg/mL chloramphenicol for pCE. *Efs* OG1RF can be selected with fusidic acid (25 µg/mL), and *Efs* CK111 can be selected with spectinomycin (1,000 µg/mL).

### Bacterial Mutagenesis and Complementation

Sequences of the primers used for mutagenesis, screening, and other plasmid construction are located in **Table S2** (Sigma Aldrich). Bacterial strains and plasmids used or generated in this study are described in **Table S1**.

Mutagenesis with the PheS counterselection system^31,32^ was used to generate deletion mutants in the *Efs* OG1RF background strain. Briefly, upstream and downstream fragments (approximately 1,000 bp each) of the gene to be deleted were amplified with Phusion (New England Biolabs) according to manufacturer instruction. Fragments were joined via crossover PCR using PrimeStar Max (Takara Bio) per manufacturer instruction. Crossover fragment ends and vector pHOU1 were digested with either BamHI, EcoRI, or XbaI (New England Biolabs) at 37 °C for 2h. T4 DNA ligase (New England Biolabs) was used to ligate the crossover fragment into pHOU1 with overnight incubation at 16°C. pHOU1 containing the flanking regions was transformed into *E. coli* EC1000, electroporated in *Efs* CK111, and conjugated into *Efs* OG1RF. After confirming the first recombination event after conjugation into *Efs* OG1RF, colonies were streaked onto minimal media agar containing p-chloro-Phe to eject the plasmid. Deletion of the gene of interest were confirmed via PCR then Sanger sequencing (Azenta) and/or whole genome sequencing.

Deletion mutants and chromosomal protein tags in *Efs* OG117 were generated using the CRISPR-Cas9 mutagenesis system^18^. For the deletion mutants, upstream and downstream flanking regions (approximately 1,000 bp) were amplified using PrimeStar HS (Takara Bio) and cloned into the vector pCE along with a 30 bp site specific spacer sequence (in the gene to be deleted) using Gibson assembly (Gibson Assembly Cloning Kit, New England Biolabs) using 1µg of fragments and plasmid at a 3:1 ratio then incubating at 50 °C for 4 hours. 2 µL of the Gibson assembly was then transformed into NEB® 5-alpha Competent *E. coli* (New England Biolabs) according to manufacturer protocol and plated onto selective LB agar containing 15 μg/mL chloramphenicol. Positive colonies were confirmed with Sanger sequencing (Azenta) and electroporated into *Efs* OG117 or its derivatives. Colonies successfully mutagenized were plated on minimal media agar containing p-chloro-Phe to eject the plasmid. Colonies containing the deletion were confirmed via Sanger sequencing (Azenta) and/or whole genome sequencing. For protein tagging, the gene encoding the fluorescent protein of interest was cloned upstream or downstream of the gene of interest with a glycine-glycine-serine linker to generate the fragment for Gibson assembly.

Complementation was carried out *in trans* from either the nisin inducible vector pMSP3535 or the constitutively expressing vector pAT392. The gene of interest was amplified by PCR with Phusion (New England Biolabs), digested with with BamHI and XbaI (New England Biolabs), and ligated into the appropriate vector with T4 DNA ligase (New England Biolabs). Recombinant plasmids were confirmed by PCR (GoTaq DNA Polymerase, Promega) and Sanger sequencing (Azenta), prior to electroporation into the strain of interest. Empty vector (for fluorescence studies, vector containing the fluorescent protein alone) were also electroporated into the strains of interest as a control.

### Minimum Inhibitory Concentrations

Minimum inhibitory concentrations for daptomycin were determined using Etest (Biomerieux). Strains were diluted to make a 0.5 McFarland standard that was then inoculated on BHI agar (Becton Dickinson) containing appropriate antibiotics (Sigma Aldrich) for selection of plasmids as described above. Etest strips were placed on the plate, incubated at 37°C, and MIC results were interpreted after 24 h.

### Quantitative Real-Time PCR

Oligonucleotides used for expression studies are described in Supplementary Table S2. Cell cultures were grown to either exponential or stationary phase in tryptic soy broth at 37°C and pelleted. RNA was extracted using the PureLink RNA Extraction Kit (Invitrogen) and contaminating DNA was removed with TurboDNase (Invitrogen). cDNA was generated using SuperScriptII (ThermoFisher) reverse transcriptase according to manufacturer instructions. 5 ng of cDNA was loaded in triplicate and qRT-PCR was used to evaluate gene expression using SsoAdvanced Universal SYBR Green Supermix (Bio-Rad) in the CFX96 Touch Real-Time PCR Detection System (Bio-Rad) as follows: 30 seconds at 95°C, 35 cycles: 15 seconds at 95°C, 30 seconds at 60°C. Relative gene expression was calculated using the Pffafl^19^ method with the gene encoding GyrB or 16S rRNA as the reference for normalization. The LinReg program was used to determine primer efficiency. t-test was used to evaluate differential gene expression between *cls1* and *cls2* for each growth condition.

### Lipid Extraction and Liquid Chromatography-Mass Spectrometry

#### Reagents

High-performance LC grade solvents (water, acetonitrile, chloroform, and methanol) and ammonium acetate (Optima LC/MS grade) were purchased from Thermo Fisher Scientific. All PC and PE lipid standards used for collisional cross section (CCS) calibration were purchased from Avanti Polar Lipids^35^.

#### Lipid extraction

Lipid extraction was conducted based on the Bligh & Dyer method as described elsewhere^36–38^. Briefly, bacteria broth was collected after cultivation in 24 hours, rinsed with 1x PBS, spined and dried with a speed-vac. 1 mL of water was then added to the pelleted and dried bacteria. The resulting suspensions were sonicated in an ice bath for 30 min to dislodge the dried pellets and homogenize the suspension. 4 mL of chilled solution of chloroform and methanol (1:2 v/v) was added to each tube, followed by 5 min of vigorous vortex and the addition of 1 mL of chilled chloroform and 1 mL of chilled water. The samples were then rigorously vortexed for 1 min and centrifuged for 10 min at 4°C and 2,000 x *g* to separate the organic and aqueous layers. The organic layers were collected to new 10 mL glass centrifuge tubes (Fisher Scientific, Waltham, MA, USA) and dried in a vacuum concentrator. The dried lipid extracts were reconstituted with 500 µL of 1:1 chloroform/methanol and stored in -80 °C. For HILIC-IM-MS analysis, 5 µL of lipid extract was transferred into LC vials, dried under nitrogen, and redissolved in 100 µL of 2:1 acetonitrile/methanol.

#### Liquid chromatography

Bacterial lipids were separated by a Waters UPLC (Waters Corp., Milford, MA, USA) as described previously^24,25^. Briefly, hydrophilic interaction liquid chromatography (HILIC) was performed with a Phenomenex Kinetex HILIC column (2.1×100 mm, 1.7 µm) maintained at 40 °C at a flow rate of 0.5 mL/min. The solvent system consisted of: A) 50% acetonitrile/50% water with 5 mM ammonium acetate, and B) 95% acetonitrile/5% water with 5 mM ammonium acetate. The gradient program was optimized as follows: 0-1 min, 100% B; 4 min, 90% B; 7-8 min, 70% B; 9-12 min, 100% B. A sample injection volume of 5 µL was used for all analyses.

#### Ion mobility-mass spectrometry

The Waters Synapt G2-XS platform was used for lipidomics analysis. Effluent from the UPLC was introduced through the electrospray ionization (ESI) source. ESI capillary voltages of +2.0 and -2.0 kV were used for positive and negative analyses, respectively. Additional ESI conditions were as follows: sampling cone, 40 V; extraction cone, 80 V; source temperature, 150 °C; desolvation temperature, 500 °C; cone gas, 10 L/hr; desolvation gas, 1000 L/hr. Mass calibration over *m/z* 50-1200 was performed with sodium formate. Calibration of ion mobility (IM) measurements was performed as previously described^39^. IM separation was performed with a traveling wave height of 40 V and velocity of 500 m/s. Data was acquired for *m/z* 50-1200 with a 1 sec scan time. Untargeted MS/MS (MS^E^) was performed in the transfer region with a collision energy ramp of 35-45 eV. Mass and drift time correction was performed post-acquisition using the leucine enkephalin lockspray signal.

#### Data analysis

Data alignment, chromatographic peaks detection, and normalization were performed in Progenesis QI (Nonlinear Dynamics). A pooled quality control sample was used as the alignment reference. The default “All Compounds” method of normalization was used to correct for variation in the total ion current amongst samples. Lipid identifications were made based on *m/z* (within 10 ppm mass accuracy), retention time and CCS with an in-house version of LipidPioneer, modified to contain the major lipid species observed in *E. faecalis*, including diacylglycerols (DGs), diglycodiacylglycerols (DGDGs), phosphatidylglycerols (PGs), cardiolipins (CLs), and LysylPGs (LPGs) with fatty acyl compositions ranging from 25:0 to 38:0 (total carbons: total degree unsaturation), and LiPydomics^24,40,41^.

Results were reported normalized to the 15:0/15:0 phosphatidylethanolamine (Avanti Polar Lipids) control for overall lipid classes by abundance or percentage of individual lipid species to the whole lipid class. t-test was used to compare. *E. facalis* OG117 to OG117Δ*liaX*. Two-way ANOVA was used to compare *E. faecalis* OG117 and the individual *cls* deletions (or *Efs* OG117Δ*liaX* and the individual *cls* deletions) and t-test was used to compare the individual *cls* deletions to each other.

### Fluorescence Microscopy

Images were adjusted for “Black Balance” per BZ-X800 Image Analysis Software with individual representative image selected for figures. All adjustments were made to whole images.

#### Anionic phospholipid microdomains/10-N-nonyl acridine orange (NAO) staining

Cells were grown in tryptic soy broth, and 1 uM NAO was added in early exponential phase. Cells were grown in the dark at 37 °C with agitation to early stationary phase prior to washing with phosphate buffered saline and adherence to poly-L-lysine coverslips. Coverslips were mounted to glass slides with ProGlass media and visualized using a Keyence BZ-X800 fluorescence microscope with filters for GFP (excitation: 470nm/emission:525nm). Quantification of at least 50 cells per strain in triplicate was done to evaluate whether regions of fluorescence intensity were septally located. One-way ANOVA was used to evaluate the differences between the strains.

#### Fluorescent tags

Proteins of interest were tagged with either GFP or mCherry. For fluorescently tagged proteins expressed in *trans*, growth in brain heart infusion (BHI) media was supplemented with appropriate antibiotics (gentamicin 150 µg/mL or erythromycin 15 µg/mL) for selection and/or induction (75-150 ng/mL nisin added after 1h of growth continuing to early stationary phase). If NAO staining was concurrent, 1 µM NAO was added in early exponential phase. Cells were grown to late stationary phase (optimal stage for concurrent expression of both fluorescently tagged proteins for visualization) in the dark at 37 °C with agitation, washed, and mounted to coverslips as described above. Visualization was performed using a Keyence BZ-X800 fluorescence microscope with filters for GFP (excitation: 470nm/emission:525nm) or Texas Red (excitation 560nm/emission: 630nm). Overlay images were produced using the BZ-X800 Analyzer software (Keyence). Representative images of protein localization/co-localization are shown.

#### Fluorescent quantification

Quantification and colocalization of fluorescence for LiaY-mCherry, NAO staining, and Cls1-ReAsH was performed using the Coloc2 plugin in ImageJ (https://imagej.nih.gov/ij/index.html). The Pearson Correlation Coefficient was calculated for 14-16 ROIs for each condition. The Fisher Transformation was used to calculate z-scores from the Pearson Correlations and determine the mean z-score for each condition. The Fisher Inverse Transformation was used to calculate the Pearson Correlation Coefficient from the mean z-score. This coefficient was used to calculate the 95% confidence interval based on the number of ROIs measured for each condition.

#### Tetracysteine-based fluorescence

Proteins of interest were tagged with a tetracysteine motif^28^ at the N-terminal domain and expressed in pMSP3535 in *E. faecalis* using erythromycin (15 µg/mL) for selection of plasmid containing clones. Cultures were grown in tryptic soy broth at 37°C for 2.5 h prior to addition of 50 ng/mL of nisin for induction. Cultures were continued for another 3.5 h prior to addition of 1 µM NAO and/or 1 µM ReAsH-EDT2 reagent (Invitrogen). Cultures were incubated for 1 h at room temperature and washed with 1X BAL wash buffer per manufacturer kit instructions. Cultures were adhered to coverslips and mounted to glass slides for visualization with a Keyence BZ-X800 with filters for GFP (excitation: 470 nm/emission:525 nm) or Texas Red (excitation 560 nm/emission: 630 nm). Representative images of protein localization/co-localization are shown.

### Bacterial two hybrid system

Oligonucleotides used for bacterial two hybrid studies are shown in **Table S1.**

Protein-protein interactions were evaluated using the Bacterial Two Hybrid System (Euromedex) according to the manufacturer instructions. Briefly, proteins of interest were tagged with catalytic adenylate cyclase domains (T18 or T25) at their N- or C-terminal domains and co-expressed in *E. coli* BTH101 (a reporter strain lacking adenylate cyclase) in Luria Bertani broth containing 50 µg/mL kanamycin, 100 µg/mL ampicillin, and 1 µM Isopropyl β-D-1-thiogalactopyranoside (IPTG) as the inducer. The strength of the interaction was determined by a β-galactosidase assay (per Euromedex instructions) after incubation at 30°C for 48-72 hours. Briefly, induced cells were permeabilized with toluene (Sigma Aldrich) and SDS (Fisher) prior to addition of 0.4% o-nitrophenol-β-galactoside (Sigma-Aldrich). Solutions were incubated at 30°C to allow for the colorimetric reaction to proceed. Reaction was stopped upon development of visible yellow color with 1M Na2CO3 (Fisher). The absorbance at OD420 nm was read using a BioTek Synergy H1 Microplate Reader. Activity units were reported normalized to cell density and reaction time.

## QUANTIFICATION AND STATISTICAL ANALYSES

Statistical details for each study can be found in the figure legends. The analyses were performed in GraphPad Prism, with the analysis type and n (number of replicates or cells counted) indicated in the figure legend for each experiment. Statistical significance is defined in each figure legend. Error bars represent standard error of the mean.

## KEY RESOURCES TABLE

Oligonucleotides (Sigma-Aldrich) used in this study can be found in Supplementary Table 2.

| Reagent | Source | Catalog Number |
| --- | --- | --- |
| Luria Bertani Broth/Agar | BD | 244620/244520 |
| Brain Heart Infusion Broth/Agar | BD | 237500/241830 |
| Tryptic Soy Broth | BD | B257543 |
| Erythromycin | Sigma-Aldrich | E5389 |
| Gentamicin | Sigma-Aldrich | G3632 |
| Daptomycin | Morris& Dickson | 650903 |
| Chloramphenicol | Sigma-Aldrich | C1919 |
| Nisin | Sigma-Aldrich | N5764 |
| Spectinomycin | Sigma-Aldrich | S4014 |
| Kanamycin | Sigma-Aldrich | K1377 |
| Ampicillin | Sigma-Aldrich | A9393 |
| IPTG | GoldBio | I2481C |
| o-nitrophenol-β-galactoside | Sigma-Aldrich | N1127 |
| Sodium carbonate | Fisher | S263-500 |
| Toluene | Fisher | T324-500 |
| 10% SDS solution | Fisher | 15553027 |
| p-chloro-phenylalanine | Sigma-Aldrich | C6506 |
| Phusion | New England BioLabs | M0530 |
| PrimeStar Max | Takara Bio | R045B |
| Gibson Assembly Cloning Kit | New England BioLabs | E5510S |
| GoTaq DNA Polymerase | Promega | M3001 |
| E-test (Daptomycin) | Biomerieux | 412323 |
| 10-N-nonyl acridine orange | Invitrogen | A1372 |
| TC-ReAsH™ II In-Cell Tetracysteine<br>Tag Detection Kit | Invitrogen | T34562 |
| Bacterial Two Hybrid Kit | Euromedex | EUK001 |
| Chloroform | FisherChemical, Optima | C2974 |
| Methanol | FisherChemical, Optima | A456-4 |
| Acetonitrile | FisherChemical, Optima | A996SK-4 |
| PE 15:0/15:0 | Avanti Polar Lipids | 850704 |

## Author Contributions

AHN, TTT, DP, AK, HV, CAA designed research. AHN, TTT, DP, VP, RZ performed research. KH, LX, RZ contributed new reagents/analytic tools. AHN, TTT, DP, KH, RZ, AK, WRM, EM, YS, LX, HV, CAA analyzed data. AHN, CAA wrote the paper.

## Acknowledgments

Project funded by NIH grant R01AI134637. We would also like to acknowledge Danielle Garsin and Kelli Palmer for advice and supplies on usage of the CRISPR-Cas9 mutagenesis system for enterococci.

