## Supplemental Data for "Molecular Basis of Cell Membrane Adaptation in Daptomycin-Resistant *Enterococcus faecalis*"

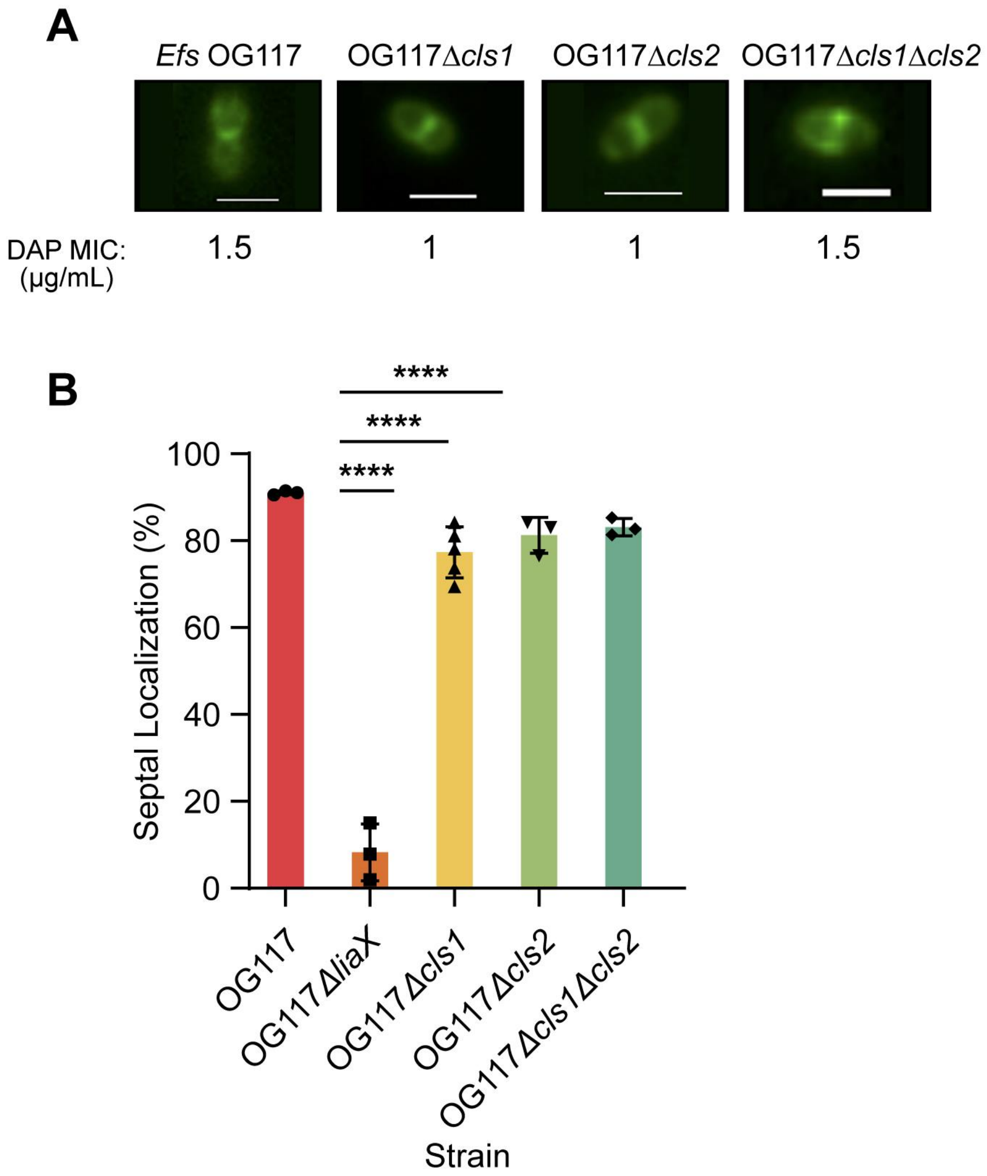


**Figure S1. Characterization of *cls* deletions in DAP-S *Efs* OG117.**

**(A)** Evaluation of anionic phospholipid localization using NAO staining in deletions of *cls1* and/or *cls2* in DAP-S *Efs* OG117.

**(B)** Quantification of septal localization of anionic phospholipid microdomains with NAO in *cls* mutant derivatives of *Efs* OG117 by counting a minimum of 50 cells per replicate (n=3-6 replicates). **p<0.01, ***p<0.001, ****p<0.0001 Whole images were adjusted for "Black Balance" per BZ-X800 Image Analysis Software with individual representative selected.

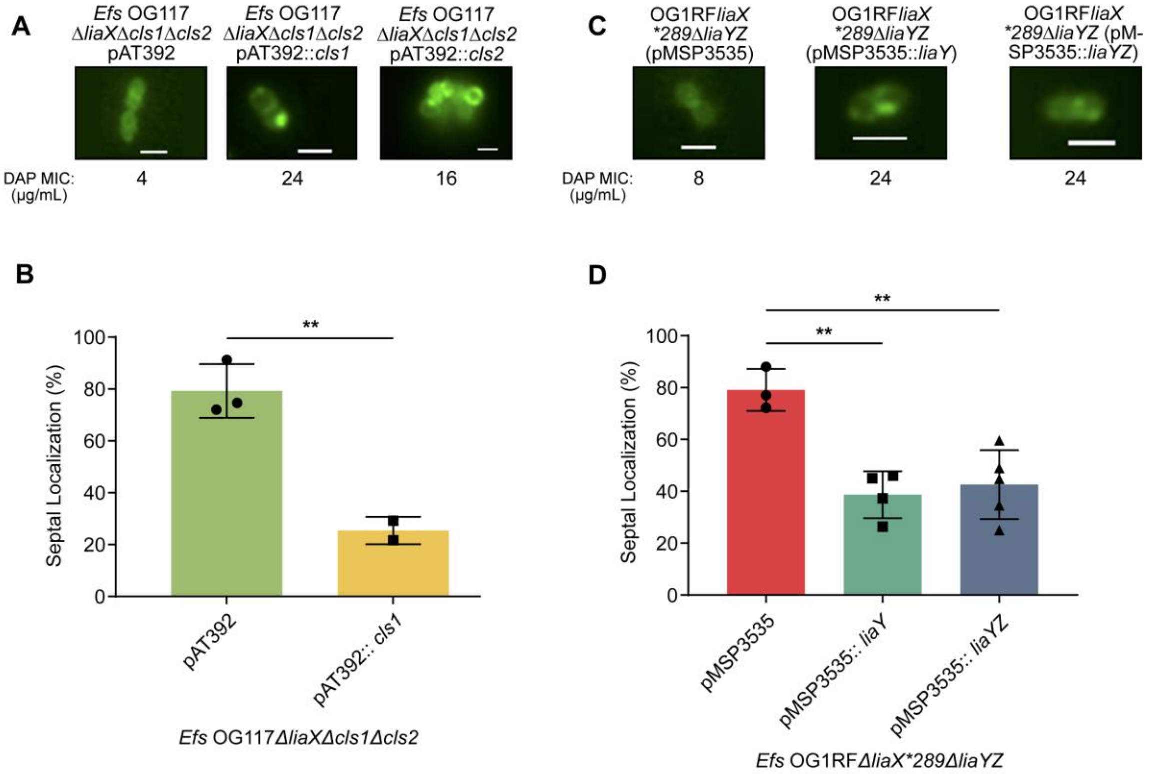


**Figure S2. Anionic phospholipid localization using NAO staining in DAP-resistant *E. faecalis* OG117Δ*liaX* and OG1RFΔ*liaX*289*Δ*liaYZ***

**(A)** *E. faecalis* OG117Δ*liaX*Δ*cls1*Δ*cls2* transformed with pAT392 and derivatives containing *cls1* or *cls2* from. Scale bar (white) at 2µm

**(B)** Quantification of septal localization of anionic phospholipid microdomains with NAO staining in *E. faecalis* OG117Δ*liaX*Δ*cls1*Δ*cls2* transformed with pAT392 and derivatives containing *cls1*. *p<0.05; **p<0.001, n=2

**(C)** *E. faecalis* OG1RFΔ*liaX*289*Δ*liaYZ* mutant transformed with pMSP3535 and derivatives containing *liaY* and *liaYZ*. Scale bar (white) at 2 µm

**(D)** Quantification of septal localization of anionic phospholipid microdomains with NAO staining in *E. faecalis* OG1RFΔ*liaX*289*Δ*liaYZ* mutant transformed with pMSP3535 and derivatives containing *liaY* and *liaYZ*. *p<0.05 **p<0.001

Whole images were adjusted for "Black Balance" per BZ-X800 Image Analysis Software with individual representative selected.

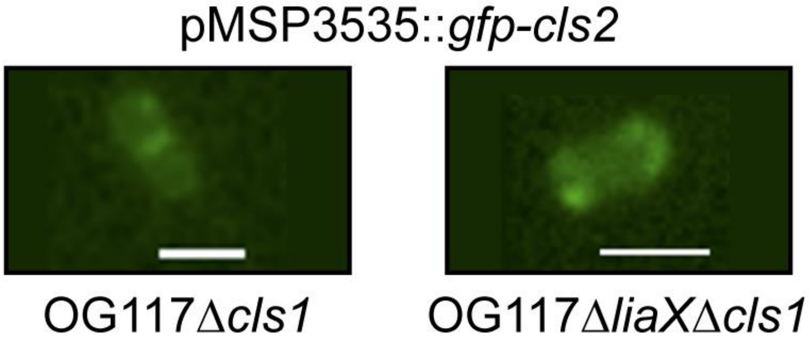


**Figure S3. Cls2 re-localizes in the absence of Cls1 in DAPS and DAP-R *E. faecalis* strains.**

Cls2 was tagged with GFP and expressed on pMSP3535 in either DAP-S *Efs* OG117Δ*cls1* or DAP-R *Efs* OG117Δ*liaX*Δ*cls1.* Introduction of *gfp-cls2* highlight the septal and non-septal localization of Cls2 in DAP-S OG117Δ*cls1* and DAP-R OG117Δ*liaX*Δ*cls1*, respectively. Scale bar (white) at 2 µm.
Whole images were adjusted for "Black Balance" per BZ-X800 Image Analysis Software with individual representative selected.

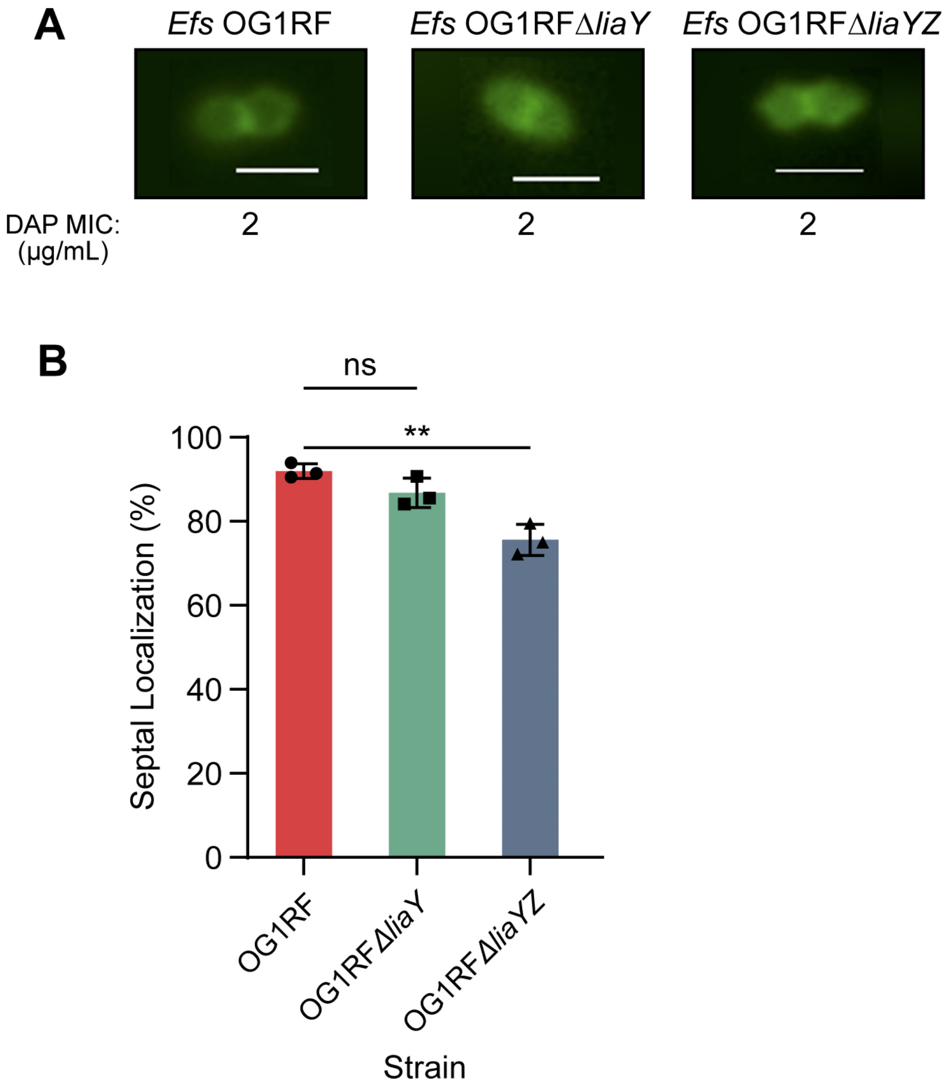


**Figure S4. Deletions of *liaYZ* in DAP-S *E. faecalis* OG1RF in which the LiaFSR is not activated**.

**(A)** NAO staining of anionic phospholipid microdomains staining with of *Efs* OG1RF, *Efs* OG1RFΔ*liaY*, and *Efs* OG1RFΔ*liaYZ* showing septal localization of microdomains without changes in DAP MIC. Scale bar (white) at 2 µm.
**(B)** Quantification of septal localization of anionic phospholipid microdomains with NAO staining in *cls* mutants. *p<0.05; **p<0.001.
Whole images were adjusted for "Black Balance" per BZ-X800 Image Analysis Software with individual representative selected.

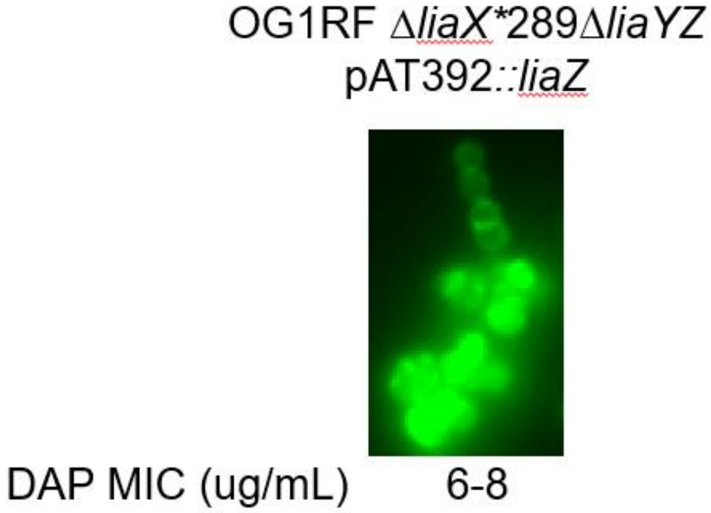


**Figure S5. *trans* expression of *liaZ* causes clumping phenotype**

Representative image of “clumping” phenotype of cells expressing *liaZ* from pAT392 and stained with 10-N-nonyl-acridine orange.
Whole images were adjusted for “Black Balance” per BZ-X800 Image Analysis Software with individual representative selected.

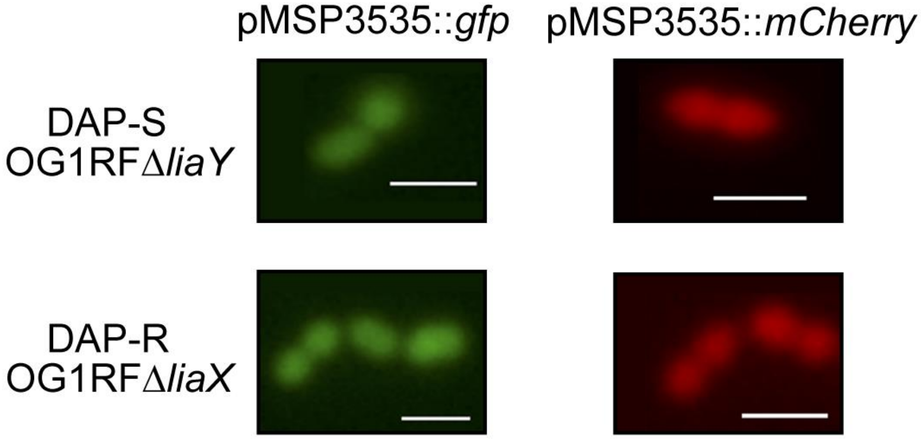


**Figure S6. Fluorescent tags do not self-aggregate.**

Genes encoding Gfp or mCherry were expressed on pMSP3535 in representative DAP-S *Efs* OG1RFΔ*liaY* and DAP-R *Efs* OG1RFΔ*liaX,* followed by visualization via fluorescence microscopy showing only a diffuse pattern with no discrete foci. Scale bar (white) at 2µm.
Whole images were adjusted for "Black Balance" per BZ-X800 Image Analysis Software with individual representative selected.

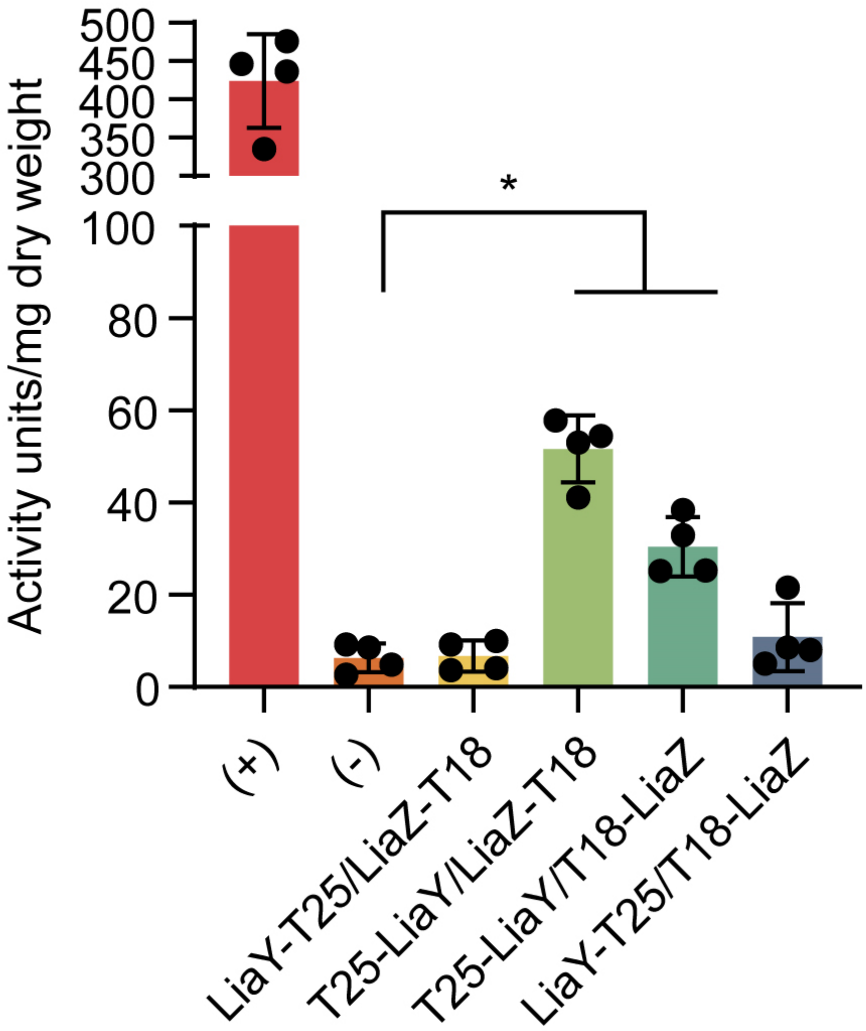


**Figure S7. LiaY interacts with LiaZ.**

LiaY-LiaZ interactions were screened via the bacterial two hybrid system. Proteins were tagged at either the N- or C-terminus, co-transformed into *E. coli* BTH101 and activity recorded via a beta galactosidase assay. A leucine zipper interaction was used as the positive control (T18-zip/T25-zip, green bar) with two non-tagged empty vectors used as negative controls (T18/T25). A positive interaction was identified between LiaY and LiaZ when LiaY was tagged at the N-terminus (T25-LiaY). Results represent the average of 3-5 experiments. *$p<0.05$

**Table S1.** ***Enterococcus faecalis*** **strains used in this study**

| Strain | DAP MIC (µg/mL) | Characteristics | Reference(s) |
|---|---|---|---|
| OG1RF | 2 | Laboratory strain | Lab strain |
| OG1RFΔ*liaX* | 12 | OG1RF containing a full deletion of *liaX* | 5 |
| OG1RF*liaX**289 | 12 | OG1RF containing a frameshift causing a truncation of predicted LiaX at amino acid position 289 | 5 |
| OG1RF*liaX**289Δ*liaZ* | 8 | OG1RF *liaX**289 with a deletion of *liaZ* | This work |
| OG1RF*liaX**289Δ*liaYZ* | 6 | OG1RF *liaX**289 with deletions of *liaY* and *liaZ* | This work |
| OG1RF*liaX**289Δ*liaYZ*(pMSP 3535) | 8 | OG1RF *liaX**289Δ*liaYZ* containing nisin-inducible pMSP3535 vector with no inserts | This work |
| OG1RF*liaX**289Δ*liaYZ* (pMSP3535::*liaY*) | 24 | OG1RF*liaX**289Δ*liaYZ* expressing *liaY* on pMSP3535 plasmid under the control of the nisin-inducible *nis* promoter. | This work |
| OG1RF*liaX**289Δ*liaYZ* pMSP3535::*liaYZ* | 24 | OG1RF*liaX**289Δ*liaYZ* expressing both *liaY* and *liaZ* on pMSP3535 plasmid under the control of the nisin-inducible *nis* promoter. | This work |
| OG1RF*liaX**289Δ*liaYZ*(pAT39 2) | 4 | OG1RF*liaX**289Δ*liaYZ c*ontaining plasmid pAT392 with no inserts. | This work |
| OG1RF*liaX**289Δ*liaYZ* pAT392::*liaY* | 8 | OG1RF*liaX**289Δ*liaYZ* expressing *liaY* on pAT392 plasmid under the control of the P2 promoter. | This work |
| OG1RF*liaX**289Δ*liaYZ* pAT392::*liaYZ* | 8 | OG1RF*liaX**289Δ*liaYZ* expressing both *liaY* and *liaZ* on plasmid pAT392 under the control of the P2 promoter. | This work |
| OG117 | 1.5 - 2 | Derivative of laboratory strain *Efs* OG1RF containing constitutively expressed *cas9* for use with CRISPR-Cas9 mutagenesis in enterococci. Representative DAP-sensitive strain | 20 |
| OG117Δ*cls1* | 1 | OG117 containing a deletion of *cls1* | This work |
| OG117Δ*cls2* | 1 | OG117 containing a deletion of *cls2* | This work |
| OG117Δ*cls1*Δ*cls2* | 1.5 | OG117 containing deletions of *cls1* and *cls2* | This work |

| | | | |
|---|---|---|---|
| OG117Δ*liaX* | 8-12 | OG117 containing a full deletion of *liaX*. Representative DAP-R strain with an activated LiaFSR system | This work |
| OG117Δ*liaX*Δ*cls1* | 8 | OG117Δ*liaX* containing a deletion of *cls1* | This work |
| OG117Δ*liaX*Δ*cls2* | 12 | OG117Δ*liaX* containing a deletion of *cls2* | This work |
| OG117Δ*liaX*Δ*cls1*Δ*cls2* | 4 | OG117Δ*liaX* containing deletions of *cls1* and *cls2* | This work |
| OG117Δ*liaX*Δ*cls1*Δ*cls2*(pAT392) | 4 | OG117Δ*liaX*Δ*cls1*Δ*cls2* containing plasmid pAT392 vector with no inserts. | This work |
| OG117Δ*liaX*Δ*cls1*Δ*cls2*(pAT392::*cls1*) | 24 | OG117Δ*liaX*Δ*cls1*Δ*cls2* containing *cls1* expressed from pAT392 vector under the control of the P2 promoter. | This work |
| OG117Δ*liaX*Δ*cls1*Δ*cls2*(pAT392::*cls2*) | 16 | OG117Δ*liaX*Δ*cls1*Δ*cls2* containing *cls2* expressed from pAT392 vector under the control of the P2 promoter. | This work |
| OG117Δ*liaX*Δ*cls1*Δ*cls2*(pMSP3535) | 4 | OG117Δ*liaX*Δ*cls1*Δ*cls2* containing nisin-inducible pMSP3535 plasmid with no inserts. | This work |
| OG117Δ*liaX*Δ*cls1*Δ*cls2*(pMSP3535::*cls1*) | 16 | OG117Δ*liaX*Δ*cls1*Δ*cls2* containing *cls1* expressed on pMSP3535 plasmid under the control of the nisin-inducible *nis* promoter. | This work |
| OG117Δ*liaX*Δ*cls1*Δ*cls2*(pMSP3535::*cls2*) | 8 | OG117Δ*liaX*Δ*cls1*Δ*cls2* containing *cls2* expressed on pMSP3535 plasmid under the control of the nisin-inducible *nis* promoter | This work |

DAP, daptomycin; MIC, minimal inhibitory concentrations.

**Table S2. Primers used in this study**

| Primer | Sequence 5' → 3' | Description (F, forward. R, reverse) |
|---|---|---|
| **Deletion of *cls1*** | | |
| FrA_*cls1*_F | atattacagctccagatccatatccttcttCAAAATTGGATCAAAAAGAA | F, lowercase: plasmid, uppercase: fragment upstream of *cls1* |
| FrA_*cls1*_R | TCATTTTGTGAAAAACAGTTTTGTCTTCCTCCTTTATTTGTTA | R, upstream of *cls1* |
| FrB_*cls1*_F | CAAATAAAGGAGGAAGACAAAACTGTTTTTCACAAAATGACG | F, downstream of *cls1* |
| FrB_*cls1*_R | gaagcgaaaaaggagaagtcggttcagaaaAAGACTGTCCACATTATGTTGC | R, lowercase: plasmid, uppercase: fragment downstream of *cls1* |
| Spacer_*cls1*_F | GTTTTGAACTTTATAGTCAAAAGATGCCGAgttttagagtcatgttgtttagaatgg | F, lowercase: plasmid, uppercase: spacer sequence within *cls1* |
| Spacer_*cls1*_R | TCGGCATCTTTTGACTATAAAGTTCAAAACtttcattgctattatacccatgtag | F, lowercase: plasmid, uppercase: spacer sequence within *cls1* |
| **Deletion of *cls2*** | | |
| FrA_*cls2*_F | agctccagatccatatccttcttACCCAAGTGATTACGATTGAC | F, lowercase: plasmid, uppercase: fragment upstream of *cls2* |
| FrA_*cls2*_R | CTGATTTATTGACGCTTACCTCCTTCTTACTTC | R, upstream of *cls2* |
| FrB_*cls2*_F | AGGAGGTAAGCGTCAATAAATCAGCAGTGAATG | F, downstream of *cls2* |
| FrB_*cls2*_R | cgaaaaaggagaagtcggttcagaaaTCTGGAATGACTTTTTCCAA | R, lowercase: plasmid, uppercase: fragment downstream of *cls2* |
| Spacer_*cls2*_F | TTACTGGCGAGATACCCATATTCGTTTGGTgttttagagtcatgttgtttagaatgg | F, lowercase: plasmid, uppercase: spacer sequence within *cls2* |
| Spacer_*cls2*_R | ACCAAACGAATATGGGTATCTCGCCAGTAAtttcattgctattatacccatgtag | F, lowercase: plasmid, uppercase: spacer sequence within *cls2* |
| **Other *cls1* constructs** | | |
| BamHI_*cls1*_pAT_F | CGGGATCCTAACAAATAAAGGAGGAAGACAATTG | F, for cloning *cls1* into pAT392, containing BamHI site |
| XbaI_*cls1*_pAT_R | CGTCTAGATTAAAGAATTGGTGAAAATAATCGTG | R, for cloning *cls1* into pAT392, containing XbaI site |
| *cls1-gfp*_FrA_F | CTCCAGATCCATATCCTTCTTGGATCATATCGGTACCAAAA | F, for construction of chromosomal insertion of GFP tag to Cls1 via CRISPR. |

| | | |
|---|---|---|
| *cls1-gfp*_FrA_R | AATTTGTAACTTCGTATATCTTGATTG | R, for construction of chromosomal insertion of GFP tag to Cls1 via CRISPR |
| *cls1-gfp*_FrA2_F | AATCAAGATATACGAAGTTACAAATTAAAC | F, for construction of chromosomal insertion of GFP tag to Cls1 via CRISPR |
| *cls1-gfp*_FrA2_R | ACTACTGCCACCAAGAATTGGTGAAAATAATCGTG | R, for construction of chromosomal insertion of GFP tag to Cls1 via CRISPR |
| *cls1-gfp*_GFP_F | CAATTCTTGGTGGCAGTAGTAAAGGAGAAGAACTTTTC | F, for construction of chromosomal insertion of GFP tag to Cls1 via CRISPR |
| *cls1-gfp*_GFP_R | GTGAAAAACAGTTTTATTTGTATAGTTCATCCATG | R, for construction of chromosomal insertion of GFP tag to Cls1 via CRISPR |
| *cls1-gfp*_FrB_F | GAACTATACAAATAAAACTGTTTTTCACAAAATGA | F, for construction of chromosomal insertion of GFP tag to Cls1 via CRISPR |
| *cls1-gfp*_FrB_R | AAGGAGAAGTCGGTTCAGAAAAAGACTGTCCACATTATGTT | R, for construction of chromosomal insertion of GFP tag to Cls1 via CRISPR |
| *cls1-gfp*_spacer_F | GCATTTGCTTCAAAGTTTAATTTGTAACTTGTTTTAGAGTCATGTTGTTTAG | F, for construction of chromosomal insertion of GFP tag to Cls1 via CRISPR |
| *cls1-gfp*_spacer_R | AAGTTACAAATTAAACTTTGAAGCAAATGCTTTCATTGCTATTATACCCATGTAG | R, for construction of chromosomal insertion of GFP tag to Cls1 via CRISPR |
| **Other *cls2* constructs** | | |
| BamHI_*cls2*_pmsp_F | CGGGATCCTAATCTTAGTATGAAGTAAGAAGGAGGTAAGCG | F, for cloning *cls2* into pMSP3535 containing BamHI site |
| XbaI_*cls2*_pmsp_R | GCTCTAGATTACAAGACTGGTGACAACAAGCG | R, for cloning *cls2* into pMSP3535 containing XbaI site |
| BamHI_*cls2*_pat_F | TCGGTACCCGGGGATCCTCTTAGTATGAAGTAAGAAGGAGGTAAGCG | F, for cloning *cls2* into pAT392 containing BamHI site |
| XbaI_*cls2*_pat_R | GCAAGGGGAATTGACTCTAGATTACAAGACTGGTGACAACAAGCG | R, for cloning *cls2* into pAT392 containing BamHI site |
| *gfp-cls2*_*cls2*_F | ACAAAGGTGGCAGTAAAATTTTTGTTTGGATTTTG | F, for cloning gfp-cls2 into pMSP3535 |
| *gfp-cls2*_*cls2*_R | AGACCGGCCTCGAGTCTAGATTACAAGACTGGTGACAAC | R, for cloning gfp-cls2 into pMSP3535 |
| *gfp-cls2*_*GFP*_F | TCTTGGGTGGCAGTAGTAAAGGAGAAGAACTTTTC | F, for cloning gfp-cls2 into pMSP3535 |
| *gfp-cls2*_*GFP*_R | AATTTTACTGCCACCTTTGTATAGTTCATCCATGC | R, for cloning gfp-cls2 into pMSP3535 |
| **Deletion of *liaY*** | | |

| | | |
|---|---|---|
| FrA_*liaY*_F | CGGGATCCGACATTGATGTGGATGATGAAA | F, fragment upstream of *liaY.* BamHI site |
| FrA_*liaY*_R | CAATCGCTGAAAATATGTCATATGTTTTCACCTCGTCTAC | R, fragment upstream of *liaY.* |
| FrB_*liaY*_F | ATGACATATTTTCAGCGATTGGTTGTTAATACACTGACAT | F, fragment downstream of *liaY.* |
| FrB_*liaY*_R | CGGAATTCAATCAAGGACGAGAACCAATC | R, fragment downstream of *liaY.* EcoRI site |
| **Deletion of *liaYZ*** | | |
| FrA_*liaYZ*_F | GCTCTAGAGACATTGATGTGGATGATGAAA | F, fragment upstream of *liaYZ.* XbaI site |
| FrA_*liaYZ*_R | ATGTTTTCACCTCATCTACTAAATTTTACACTAGAAAGGC | R, fragment upstream of *liaYZ.* |
| FrB_*liaYZ*_F | AGTAGATGAGGTGAAAACATAAAAAGCCTGAAGAACGTAGCA | F, fragment downstream of *liaYZ.* |
| FrB_*liaYZ*_R | CGGGATCCATCGGCCGCGCTACAATCGC | R, fragment downstream of *liaYZ.* BamHI site |
| **Other *liaY* constructs** | | |
| BamHI_*liaY*_F | CGGGATCCACGAGGTGAAAACATATGAAAAG | F, upstream of *liaY* start codon BamHI site |
| XbaI_*liaY*_R | GCTCTAGATTAAAAGTCACTCCATTCGTCATC | R, 3’ end of *liaY* including stop codon. XbaI site |
| *liaY-mCherry_liaY*_F | CAGGAGACTCTGCATGGATCCTAGTAGACGAGGTGAAAACATATG | F, for cloning of liaY-mCherry into pMSP3535, contains BamHI site |
| *liaY-mCherry_liaY*_R | CTTGGAAACGCTTCCGCCACAACATCCTGGACAACAGC | R, for cloning of liaY-mCherry into pMSP3535 |
| *liaY-mCherry_mCherry*_F | GTGGCGGAAGCGTTTCCAAGGGCGAGGAGG | F, for cloning of liaY-mCherry into pMSP3535 |
| *liaY-mCherry_mCherry*_R | AGACCGGCCTCGAGTCTAGATTATTTGTACAGCTCATCCATGCC | R, for cloning of liaY-mCherry into pMSP3535, contains XbaI site |
| *liaY-mCherry*_Q5_F | GTTTCCAAGGGCGAGGAG | F, generation of pMSP3535 mCherry control through Q5 mutagenesis |
| *liaY-mCherry*_Q5_R | CATATGTTTTCACCTCGTCTAC | R, generation of pMSP3535 mCherry control through Q5 mutagenesis |

| | | |
|---|---|---|
| **Complementation of *liaYZ*** | | |
| BamHI_*liaYZ*_F | CGGGATCCACGAGGTGAAAACATATGAAAAG | F, upstream of *liaY* start codon BamHI site |
| XbaI_*liaYZ*_R | GCTCTAGATTATTCATATTTCGATAAATTGTG | R, 3' end of *liaZ* including stop codon. XbaI site |
| **Bacterial Two Hybrid Studies** | | |
| *cls1*_F | CGCGGATCCGATTCTTAGCGTCTTAACAGT | F, for cloning *cls1* into pKT25/pKNT25 or pUT18/pUT18C |
| *cls1*_R | CCGGAATTCAGAATTGGTGAAAATAATCGT | R, for cloning *cls1* into pKNT25/pUT18 |
| *cls1*_R2 | CCGGAATTCTTTAAAGAATTGGTGAAAAT | R, for cloning *cls1* into pKT25/pUT18C, contains stop codon |
| *cls2*_F | CGCGGATCCGAAAATTTTTGTTTGGATTTT | F, for cloning *cls2* into pKT25/pKNT25 or pUT18/pUT18C |
| *cls2*_R | CCGGAATTCAAGACTGGTGACAACAAGC | R, for cloning *cls2* into pKNT25/pUT18 |
| *cls2*_R2 | CCGGAATTCATTACAAGACTGGTGACAACA | R, for cloning *cls2* into pKT25/pUT18C, contains stop codon |
| *liaY*_F | CGCGGATCCGAAAAGAAAACTAACAAAATCTCCG | F, for cloning *liaY* into pKT25/pKNT25 or pUT18/pUT18C |
| *liaY*_R | CCGGAATTCAAGTCACTCCATTCGTCATC | R, for cloning *liaY* into pKNT25/pUT18 |
| *liaY*_R2 | CCGGAATTCATTAAAAGTCACTCCATTCGTCAT | R, for cloning *liaY* into pKT25/pUT18C, contains stop codon |
| *liaZ*_F | CGCGGATCCGACATATTTTCAGCGATTGGT | F, for cloning *liaZ* into pKT25/pKNT25 or pUT18/pUT18C |
| *liaZ*_R | CCGGAATTCTCATATTTCGATAAATTGTGAT | R, for cloning *liaZ* into pKNT25/pUT18 |
| *liaZ*_R2 | CCGGAATTCTTTTATTCATATTTCGATAAATTGTG | R, for cloning *liaZ* into pKT25/pUT18C, contains stop codon |
| **qRT-PCR** | | |
| *16S*_F | GGAGACTTGAGTGCAGAAGA | F, Housekeeping gene, 16S rRNA of OG1RF |
| *16S*_R | CGTCAGTTACAGACCAGAGAG | R, Housekeeping gene, 16S rRNA of OG1RF |
| *gyrB*_F | AAAAGGCATGTTGGCTTCAAA | F, Housekeeping gene, *gyrB* of OG1RF |

| | | |
|---|---|---|
| *gyrB*_R | GCTTCCCTGGCAAGTTGCTA | R, Housekeeping gene, *gyrB* of OG1RF |
| *cls1*_F | GGTGCCAGAGGCAAATACAT | F, to amplify *cls1* for qRTPCR |
| *cls1*_R | AATACGGCGTTTGAATCCAG | R, to amplify *cls1* for qRTPCR |
| *cls2*_F | GTTGTTAGTGGCGGTTCGTT | F, to amplify *cls2* for qRTPCR |
| *cls2*_R | TTAATGCCGCGTTTGTGTAA | R, to amplify *cls2* for qRTPCR |
